# Fgfr2b signaling is essential for the maintenance of the alveolar epithelial type 2 lineage during lung homeostasis in mice

**DOI:** 10.1101/2022.01.26.477823

**Authors:** Negah Ahmadvand, Arun Lingampally, Farhad Khosravi, Ivonne Vazquez-Armendariz, Stefano Rivetti, Jochen Wilhelm, Susanne Herold, Guillermo Barreto, Janine Koepke, Christos Samakovlis, Gianni Carraro, Jin-San Zhang, Denise Al Alam, Saverio Bellusci

**Author notes:** Correspondence to: Saverio Bellusci. Author’s contributions: N.A designed the study, carried out the experiments, analyzed the data and wrote the manuscript. A.L. contributed to the experiments and quantification analysis. F.K contributed to performing experiments, data analysis and writing of the manuscript. A.I.V.A contributed to the experiments and quantification analysis. S.R. contributed to the experiments and provided feedback in the writing of the manuscript. J.W. and J.K. contributed to the experiments and data analysis. S.H., G.B., J.Z., C.S. and D.A.A. provided feedback, helped shape the research, discussed the results, and contributed to the final manuscript. S.B designed the project, regularly monitored the generated results, interpreted the results and wrote the manuscript in coordination with N.A. All authors reviewed the results and contributed to the final manuscript.

## Abstract

Fibroblast growth factor receptor 2b (Fgfr2b) signaling is essential throughout lung development to form the alveolar epithelial lineage. However, its role in alveolar epithelial type 2 cells (AT2s) homeostasis was recently considered dispensable. SftpcCreERT2; *tdTomato^flox/flox^* mice were used to delete *Fgfr2b* expression in cells belonging to the AT2 lineage, which contains mature AT2s and a novel SftpcLow lineage-traced population called “injury activated alveolar progenitors” or IAAPs. Upon continuous tamoxifen exposure for either one or two weeks to delete *Fgfr2b*, a shrinking of the AT2s is observed. Mature AT2s exit the cell cycle, undergo apoptosis and fail to form alveolospheres in vitro. However, the lung morphometry appears normal, suggesting the involvement of compensatory mechanisms. In mutant lungs, IAAPs which escaped *Fgfr2b* deletion expand, display enhanced alveolosphere formation in vitro and increase drastically their AT2 signature suggesting differentiation towards mature AT2s. Interestingly, a significant increase in AT2s and decrease in IAPPs occurs after a one-week tamoxifen exposure followed by an eight-week chase period. While mature AT2s partially recover their alveolosphere formation capabilities, the IAAPs no longer display this property. Single-cell RNA seq analysis confirms that AT2s and IAAPs represent stable and distinct cell populations and recapitulate some of their characteristics observed in vivo. Our results underscore the essential role played by Fgfr2b signaling in the maintenance of the AT2 lineage in the adult lung and suggest that the IAAPs could represent a new population of AT2 progenitors.

## INTRODUCTION

The fibroblast growth factor (Fgf) family is made of 22 members. Fgfs can either act in a paracrine, endocrine or intracellular fashion. The Fgfs acting through a paracrine mechanism elicit their signaling through fibroblast growth factor receptors (Fgfr) and heparin-sulfate proteoglycans. The endocrine Fgfs signal through Fgfr with the Klotho family of proteins as co-receptors, and the intracellular Fgfs display Fgfr independent signaling [1–3]. The paracrine Fgfs contain Fgf3, 7, 10, 22 and interact mainly with Fgfr2b [4]. Among the paracrine Fgfs, Fgf10 takes center stage for its non-redundant role during development, homeostasis and repair after injury [5–7]. During the pseudoglandular stage of lung development, Fgf10 is expressed dynamically in the mesenchyme in association with the newly formed epithelial buds [8]. Genetic inactivation of *Fgf10* or its receptor *Fgfr2b* leads to a lung displaying the rudimentary primary bronchi but lacking further ramifications [9–11]. Using an inducible dominant-negative Fgfr2b approach, we characterized both primary transcriptional targets and the main biological activities associated with Fgfr2b signaling. At E12.5, Fgf10 signaling essentially regulates adherens junction and basement membrane organization. Fgf10 acts primarily through beta-catenin signaling and maintains the expression of Sox9, a transcription factor essential for alveolar progenitor differentiation, in the distal epithelium [12]. At E14.5, Fgfr2b signaling controls proliferation of the alveolar epithelial progenitors, and the identified primary transcriptional targets support both overlapping and distinct biological activities compared to E12.5 [13]. At E16.5, Fgfr2b signaling prevents the differentiation of AT2 progenitors towards the AT1 fate (Jones and Bellusci, unpublished data). Such function is conserved during the alveolar phase of lung development in mice [14].

Fgf10 also plays a vital role during the repair process. For example, *Fgf10* deletion in peribronchial mesenchymal cells leads to impaired repair following injury to the bronchial epithelium using naphthalene [15, 16]. On the other hand, overexpression of *Fgf10* reduces the severity of lung fibrosis in bleomycin-induced mice [17]. Despite these diverse biological activities during development and repair after injury, Fgfr2b signaling in AT2s has been deemed dispensable during homeostasis [14, 18]. Notably, the respective function of Fgfr2b signaling in our recently described, lineage-traced, AT2 subpopulations has not been defined [19].

Previous studies using the 3D matrigel-based alveolosphere assay in vitro and following diphtheria toxin (DTA)-based genetic deletion of lineage-labeled SftpcPos cells using *S*ftpcCreERT2/^+^*; Rosa26^LSL-DTA/LSL-tdTomato^* mice in vitro demonstrated the relevance of AT2s as stem cells for the respiratory epithelium [20, 21]. However, in both assays, the self-renewal capability is present only in a subpopulation of lineage-labeled SftpcPos AT2s as only 1-2% of the cultured FACS-isolated lineage-labeled AT2s generated alveolospheres [20]. AT2 stem cells reside in a stromal niche made of lipofibroblasts (LIFs) [5, 20, 22–24]. Some of these LIFs express Fgf10, which acts on the LIFs themselves via Fgfr1b and Fgfr2b to maintain their differentiation [5, 22]. Given the role of Fgf10^Pos^-LIFs in maintaining AT2 stem cell proliferation [24], we propose that Fgf10 signaling to AT2s via Fgfr2b could be instrumental for the maintenance of the AT2 stem cell characteristics.

Using the *S*ftpcCreERT2/^+^*; tdtomato^flox/+^* mice, we previously reported the existence of two distinct AT2 subpopulations called AT2-Tom^Low^ (aka injury-activated alveolar progenitors (IAAPs)) and AT2-Tom^High^ (aka AT2s) [19]. IAAPs express a lower level of *Fgfr2b* and *Etv5,* indicating minor Fgfr2b signaling in these cells and a low level of AT2 differentiation markers *Sftpc*, *Sftpb*, *S*ftpa1. On the other hand, AT2s show high *Sftpc*, *Sftpb*, *S*ftpa1 and significant activation of Fgfr2 signaling illustrated by the high level of *Fgfr2b* and *Etv5* expression. ATAC-seq analysis indicates these two subpopulations are distinct. Upon pneumonectomy, the number of IAAPs but not AT2s increases and IAAPs display increased expression of *Fgfr2b*, *Etv5*, *Sftpc*, *Ccnd1* and *Ccnd2* compared to sham. Therefore, our previous work suggested that IAAPs represent quiescent, immature AT2-progenitor cells in mice that could proliferate and differentiate into mature AT2s upon pneumonectomy.

This study analyzed the impact of *Fgfr2b* deletion on AT2s and IAAPs during homeostasis. We have used *S*ftpcCreERT2*; tdTomato^flox/ flox^* mice to lineage-trace AT2s and IAAPs and delete *Fgfr2b* expression in these subpopulations. In addition, flow cytometry, qPCR, ATAC-seq, gene arrays, scRNA-seq, immunofluorescence, alveolosphere assays and lung morphometry were carried out. Contrary to previous studies, our results indicate an essential role for Fgfr2b signaling in IAAPs and AT2s during homeostasis and unravel the unexpected behavior of IAAPs, likely representing a novel AT2 subpopulation with regenerative capabilities.

## MATERIALS AND METHODS

### Animal experiments

All animals were housed under specific pathogen-free (SPF) conditions with free access to food and water. Genetically modified mice including *S*ftpctm1(cre/ERT2,rtTA)Hap (stock number 007905), *Fgfr2^tm1Dsn^* (*Fgfr2-IIIb^flox^*) (gift from C. Dickson, [4](De Moerlooze et al., 2000)) and the Cre reporter line *tdTomato^flox^* (B6;129S6-Gt(ROSA)26Sor^tm9(CAG-tdTomato)Hze^/J (stock number 007909) were purchased from Jackson Laboratory (Bar Harbor/ME, USA). 8-16-weeks-old mice were treated with tamoxifen-containing water (1 mg/ml) (T5648, Sigma-Aldrich, Darmstadt/Germany) to induce Cre recombinase activity. All animal studies were performed according to protocols approved by the Animal Ethics Committee of the Regierungspraesidium Giessen (permit numbers: G7/2017–No.844-GP and G11/2019–No. 931-GP).

### Lung dissociation and FACS

Adult mice were sacrificed, and lungs were perfused with 5 ml PBS through the right ventricle. Next, lungs were inflated via the trachea with dispase and kept in dispase (Coning, NY, USA) and Collagenase Type IV at 37°C for 40 min with frequent agitation. To obtain single-cell suspensions, the digested tissue was then passed serially through 100-, 70- and 40-μm cell strainers (BD Biosciences). First, red blood cells (RBC) were eliminated using RBC lysis buffer (Sigma Aldrich), according to the manufacturer’s protocol. Next, cells were pelleted, resuspended in FACS buffer (0.1% sodium azide, 5% fetal calf serum (FCS), 0,05% in PBS) and stained with antibodies: anti-EpCAM (APC-Cy7-conjugated, Biolegend,1:50), CD49F (APC-conjugated, Biolegend,1:50), anti-PDPN (FITC-conjugated, Biolegend, 1:20) and anti-CD274 (unconjugated, Thermo Fisher, 1:100) antibodies for 20 minutes on ice in the dark, followed by washing. Then, the cells were stained for goat anti-rabbit secondary antibody Alexa flour 488 (Invitrogen,1:500) for 20 minutes on ice in the dark, followed by washing. Next, cells were washed and stained with SYTOX (Invitrogen), a live/dead cell stain according to the manufacturer’s instructions. Finally, flow cytometry data acquisition and cell sorting were carried out using FACSAria III cell sorter (BD Biosciences, San Jose/CA). Data were analyzed using FlowJo software version X (FlowJo, LLC).

### Hematoxylin and eosin staining

Mouse lung tissues were fixed using 4% PFA followed by embedding in paraffin. Paraffin blocks were sectioned into 5-μm-thick slices and placed on glass slides. Following deparaffinization, lung sections were stained with hematoxylin (Roth) for 2 min, washed with running tap water for 10 min and then stained with eosin (Thermo Fisher Scientific) for 2 min.

### RNA extraction and quantitative real-time PCR

Following lysis of FACS-isolated cells from mouse or human lungs in RLT plus, RNA was extracted using an RNeasy Plus Micro kit (Qiagen), and cDNA synthesis was carried out using QuantiTect reverse transcription kit (Qiagen) according to the manufacturer’s instruction. After that, selected primers (Table 1,2) were designed via NCBI’s Primer-BLAST option (https://www.ncbi.nlm.nih.gov/tools/primer-blast/) (for primer sequence see supplementary table). Then, quantitative real time polymerase chain reaction (qPCR) was performed using PowerUp SYBR Green Master Mix kit according to the manufacturer’s protocol (Applied Biosystems) and LightCycler 480 II machine (Roche Applied Science). *Hypoxanthine guanine phosphoribosyltransferase* (*Hprt*) was used as a mouse reference gene. Data were presented as mean expression relative to *Hprt* and assembled using the GraphPad Prism software (GraphPad Software, La Jolla/CA). Statistical analyses were performed utilizing two tailed-paired Student t-test, and results were significant when p < 0.05.

**Table 1:**
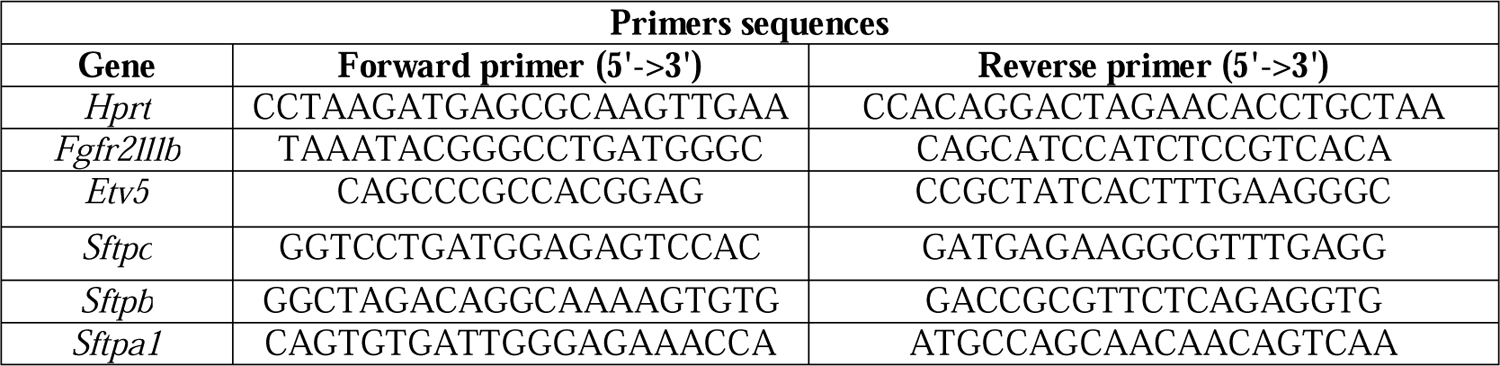

**Table 2:**
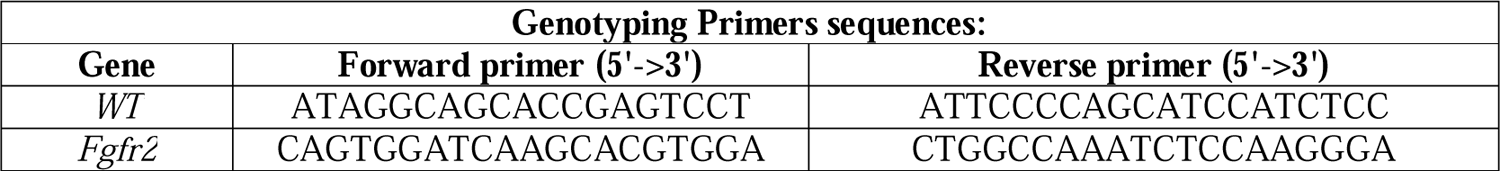

### Immunofluorescent Staining

After lung perfusion with PBS through the right ventricle, isolated lungs were fixed with 4% paraformaldehyde. Afterwards, tissues were embedded in paraffin and sectioned at 5 μm thickness. Following deparaffinization, slides were blocked with 3% bovine serum albumin (BSA) (Jackson Immunoresearch Laboratories) in PBS for 1 hour at RT. Next, immunofluorescence staining was performed using overnight incubation with polyclonal anti-Prosurfactant Protein C (ProSP-C) (Merck/Millipore/Sigma Aldrich, 1:500) followed by staining with polyclonal secondary antibody Goat anti-rabbit Alexa fluor 488 (Invitrogen,1:500). Finally, slides were mounted with ProLong Gold Antifade Reagent containing DAPI (Molecular Probes). Proliferation was assessed using the Click-iT EdU Imaging Kit (Invitrogen, Schwerte, Germany) according to the manufacturer’s instructions. For the EdU experiments, EdU was injected (i.p.) two hours before mice were sacrificed (Dosage: 0.005 mg EdU/g mouse weight). Apoptosis was assessed on paraffin sections via the TdT-mediated dUTP Nick-End Labelling (TUNEL) assay using the DeadEnd Fluorometric TUNEL System (Promega, Walldorf, Germany) according to the manufacturer’s instructions. Apoptosis was quantified by determining the ratio of TUNEL-positive cells to total cells in each region of interest. Multiple images (nL>L8) were acquired and quantified. For each experiment, sections from at least four independent lungs were analyzed.

### Alveolosphere assay

Sorted epithelial cells (IAAPs/Tom^Low^ and AT2s/Tom^High^) from [*S*ftpcCreERT2/^+^*; tdTom^flox/flox^*] mice and resident mesenchymal cells from C57BL/6J mice (Epcam^Neg^, Cd31^Neg^, Cd45^Neg^, Sca1^Pos^) were centrifuged and resuspended separately in cell culture medium (Dulbecco’s Modified Eagle Medium, Life Technologies). First, 1×10^4^ epithelial cells in 25 μL media and 2×10^4^ mesenchymal cells in 25 μL media per insert (12 mm cell culture inserts with 0.4 µm membrane Millipore) were prepared. Next, mesenchymal and epithelial cell suspensions were mixed, followed by the addition of cold Matrigel® growth factor-reduced Matrigel (Corning) at a 1:1 dilution resulting in 100 μL final volume per insert. Then, Matrigel cell suspensions were placed on the top of the filter membrane of the insert and incubated at 37°C for L of the medium was transferred to each well. Finally, cells were incubated in air-liquid interface conditions at 37°C with 5% CO2 for two weeks. Media were changed 3 times per week.

### Microarray

Purified total RNA was amplified using the Ovation PicoSL WTA System V2 kit (NuGEN Technologies). Per sample, 2 µg amplified cDNA was Cy5-labeled using the SureTag DNA labeling kit (Agilent). Hybridization to 8×60K 60mer oligonucleotide spotted microarray slides (Human Mouse Genome, Agilent Technologies, design ID 074809) and subsequent washing and drying of the slides were performed following the Agilent hybridization protocol in Agilent hybridization chambers, with following modifications: 3 µg of the labeled cDNA were hybridized for 22 hours at 65°C. The cDNA was not fragmented before hybridization.

The dried slides were scanned at 2 µm/pixel resolution using the InnoScan is900 (Innopsys). Image analysis was performed with Mapix 6.5.0 software, and calculated values for all spots were saved as GenePix results files. Stored data were evaluated using the R software and the limma package^28^ from BioConductor. Log2 mean spot signals were taken for further analysis. Data were background corrected using the NormExp procedure on the negative control spots and quantile-normalized ^28,29^ before averaging. Log2 signals of replicate spots were averaged, and from several different probes addressing the same gene, only the probe with the highest average signal was used. Genes were ranked for differential expression using a moderated t-statistic. Finally, pathway analyses were done using gene set tests on the ranks of the t-values. Pathways were taken from the KEGG database (http://www.genome.jp/kegg/pathway.html).

### ATAC-seq

25.000 FACS-sorted cells were collected and used for ATAC Library preparation using Tn5 Transposase from Nextera DNA Sample Preparation Kit (Illumina). The cell pellet was resuspended in 50 µl Lysis/Transposition reaction (12.5 µl THS-TD-Buffer, 2.5 µl Tn5, 5 µl 0.1% Digitonin, and 30 µl water) and incubated at 37°C for 30 min with occasional snap mixing. Following purification of the DNA, fragments were done by Min Elute PCR Purification Kit (Qiagen). Amplification of the Library together with Indexing Primers was performed as described. Libraries were mixed in equimolar ratios and sequenced on the NextSeq500 platform using V2 chemistry. Trimmomatic version 0.38 was employed to trim reads after a quality drop below a mean of Q15 in a window of 5 nucleotides. Only reads longer than 15 nucleotides were cleared for further analyses. Trimmed and filtered reads were aligned versus (vs) the mouse genome version mm10 (GRCm38) using STAR 2.6.1d with the parameters “--outFilterMismatchNoverLmax 0.1 --outFilterMatchNmin 20 -- alignIntronMax 1 --alignSJDBoverhangMin 999 --outFilterMultimapNmax 1 -- alignEndsProtrude 10 ConcordantPair” and retaining unique alignments to exclude reads of uncertain origin. Reads were further deduplicated using Picard 2.18.16 (Picard: A set of tools (in Java) for working with next-generation sequencing data in the BAM format) to mitigate PCR artefacts leading to multiple copies of the same original fragment. Reads aligning to the mitochondrial chromosome were removed. The Macs2 peak caller version 2.1.2 was employed to accommodate the range of peak widths typically expected for ATAC-seq^30^. The minimum qvalue was set to −4, and FDR was changed to 0.0001. Peaks overlapping ENCODE blacklisted regions (known misassemblies, satellite repeats) were excluded.

To be able to compare peaks in different samples to assess reproducibility, the resulting lists of significant peaks were overlapped and unified to represent identical regions. Sample counts for union peaks were produced using bigWigAverageOverBed (UCSC Toolkit) and normalized with DESeq2 1.18.1 to compensate for differences in sequencing depth, library composition, and ATAC-seq efficiency. Peaks were annotated with the promoter of the nearest gene in range (TSS +-5000 nt) based on reference data of GENCODE vM15.

### scRNA-seq

Single-cell suspensions were processed using the 10x Genomics Single Cell 3′ v3 RNA-seq kit. Gene expression libraries were prepared according to the manufacturer’s protocol. In addition, MULTI-seq barcode libraries were retrieved from the samples and libraries were prepared independently.

### Sequencing and processing of raw sequencing reads

Sequencing was done on Nextseq2000, and raw reads were aligned against the mouse genome (mm10, ensemble assembly 104) and mapped and counted by StarSolo (Dobin et al., doi: 10.1093/bioinformatics/bts635) followed by secondary analysis in Annotated Data Format. Pre-processed counts were analyzed using Scanpy (Wolf et al., doi: 10.1186/s<u>13059-017-1382</u>-0). Basic cell quality control was conducted by considering the number of detected genes and mitochondrial content. Cells expressing less than 300 genes or having a mitochondrial content of more than 20% were removed from the analysis. Further, we filtered genes if detected in less than 30 cells (<3%). Raw counts per cell were normalized to the median count over all cells and transformed into log space to stabilize the variance. We initially reduced the dimensionality of the dataset using PCA, retaining 50 principal components. Subsequent steps, like low-dimensional UMAP embedding (McInnes & Healy, https://arxiv.org/abs/1802.03426) and cell clustering via community detection (Traag et al., https://arxiv.org/abs/1810.08473), were based on the initial PCA. Final data visualization was done by the cellxgene package. The raw data have been deposited in GEO accession number GSE (pending).

## RESULTS

### IAAP and AT2 subpopulations respond differently to *Fgfr2b* deletion

We recently isolated two AT2 lineage-labeled SftpcPos cells (called IAAPs and mature AT2) in the mouse adult lung based on differential levels of Tomato expression [19]. A common assumption in the field is that the *Rosa26* locus is ubiquitous (expressed in all the cells) and homogeneous (expressed at the same level in all the cells of the body). Against these assumptions, the initial analysis of the *Rosa26*-LacZ mice already established that LacZ expression was not uniform throughout the embryo [25]. In addition, LacZ expression was also described to be heterogeneous in the adult mouse (Jackson lab (129-*Gt(ROSA)26Sor*/J Stock No: 002292). Interestingly, IAAPs and AT2s are observed regardless of whether *tdTomato^flox/flox^* or *tdTomato^flox/+^* reporter mice were used [19]. These results support the conclusion that it is primarily the level of tomato expression from the *Rosa26* promoter and not the efficiency of recombination of the *LoxP-STOP-LoxP-tomato* cassette downstream of the *Rosa26* promoter, which is different in these two subpopulations.

To unravel the function of Fgfr2b signaling in the AT2 lineage, [*S*ftpcCreERT2/^+^*; Fgfr2b^flox/flox^; tdTom^flox/flox^*] (*Fgfr2b-*cKO) mice, called experimental (Exp.) group, were initially treated with tamoxifen water for 7 days (Figure 1a, b). [*S*ftpcCreERT2/^+^*; Fgfr2b^+/+^; tdTom^flox/flox^*] mice undergoing the same treatment were used as controls (Ctrl).

**Fig. 1.**
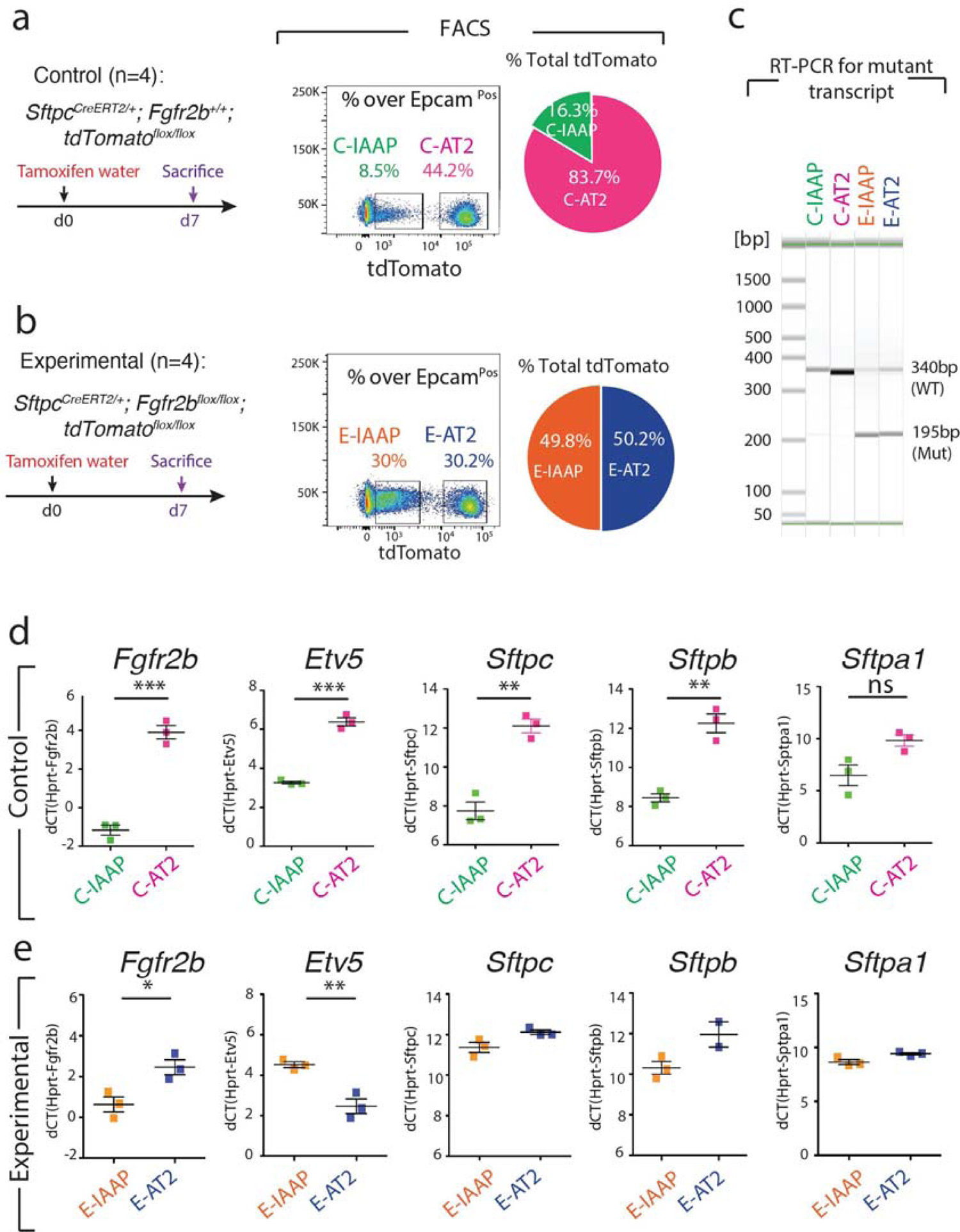
IAAP and AT2 subpopulations respond differently to *Fgfr2b* deletion. **a)** Timeline of tamoxifen treatment of *S*ftpcCreERT2/^+^*; Fgfr2b^+/+^; tdTom^flox/flox^* mice (n=4). Flow cytometry plots represent the detection of C-IAAP, C-AT2 subpopulations based on the tdTomato level in Ctrl lungs. The pie chart shows the percentage of C-IAAPs, C-AT2s in total tdTomato positive cells in the Ctrl group. **b)** Timeline of tamoxifen treatment of *S*ftpcCreERT2/^+^*; Fgfr2b ^flox/flox^; tdTom^flox/flox^* mice (n=4). Flow cytometry plots represent the detection of E-IAAPs, E-AT2s based on the tdTomato level in Exp. lungs. The pie chart shows the percentage of E-IAAPs, E-AT2s in total tdTomato positive cells in the Exp. group. **c)** RT-PCR for detecting the *Fgfr2b* mutant transcript in FACS-based sorted C-IAAPs, C-AT2s, E-IAAPs and E-AT2s. Wild type and mutant forms are detected by the size of 340bp and 195bp, respectively. **d)** qPCR analysis of FACS-based sorted C-IAAPs and C-AT2s.**e)** qPCR analysis of FACS-based sorted E-IAAPs and E-AT2s. Data are presented as mean values ± SEM. *p < 0.05, **p < 0.01, ***p < 0.001

Flow cytometry analysis on harvested lungs underpinned the presence of IAAP (AT2-Tom^Low^) and AT2 (AT2-Tom^High^) populations. We named the IAAP or AT2 cells from the Ctrl animals, C-IAAPs and C-AT2s and the IAAP or AT2 cells from the Exp. animals, E-IAAPs and E-AT2s. In Ctrl lungs, we observed an average of 9.93% +/- 1.13% (n=4) C-IAAPs and 44.75 % +/- 1.22% (n=4) of C-AT2s (of total Epcam^Pos^) as previously described (Figure 1a) [19]. In Exp. lungs, we found that E-IAAPs represented 27.05% (27.05% ± 1.83%, n=4) of the overall Epcam^Pos^ cells, and the E-AT2s represented 25.70 % (25.70% ± 2.45%, n=4) of the overall Epcam^Pos^ cells (Figure 1b). The decrease in the number of AT2 cells in Exp. vs Ctrl lungs (25.70% vs 44.75 %, respectively) indicates that the Fgfr2b pathway is critical for maintaining AT2s.

Interestingly, a concomitant increase in the percentage of IAAPs in Exp. vs Ctrl was observed (27.05% vs 9.93%), suggesting that IAAPs, previously quiescent, are becoming active and proliferative.

Next, we examined whether *Fgfr2b* is haplo-sufficient in alveolar epithelial lineage by investigating the consequences of losing a single copy of *Fgf2b* on the percentage of AT2s and IAAPs. Using *Fgfr2b* hets mice and Ctrl *Fgf2b^+/+^* (WT) mice, we carried out the previously described FACS-based approach to isolate AT2s and IAAPs [19]. No difference in the percentage of IAAPs and AT2s in WT (*Fgfr2b^+/+^*) vs *Fgfr2b^+/-^* hets could be detected, indicating that *Fgfr2b* is haplosufficient in Sftpc-expressing cells. (Figure S1).

To investigate the efficiency of recombination in the IAAPs and AT2s in Exp. (*S*ftpcCreERT2/^+^*; dTomato^flox/flox^; Fgfr2b^flox/flox^*) (n=3) vs Ctrl lungs (*S*ftpcCreERT2/^+^*; dTomato^flox/flox^; Fgfr2b^+/+^*) (n=2) (Figure S2), we analyzed the lungs 36 hours after a single dose of Tam IP. FACS analysis was carried out to quantify the abundance of IAAPs and AT2s (out of Epcam) in Ctrl and Exp. lungs. We chose the early 36-hour time point to analyze the recombination events before the possible onset of a phenotype linked to *Fgfr2b* deletion, impacting the IAAPs/AT2s ratio. As a quality control, we found a similar percentage of Epcam positive cells over total cells in Exp. vs Ctrl (23.4% vs 21.1%, respectively). Next, we analyzed the percentage of AT2s, which at later time points (at day 7 and 14 on continuous Tam water) is significantly decreased in Exp. vs Ctrl lungs (Figure 1b). As we used *tdTomato^flox/flox^* mice, we observed two peaks for the AT2s corresponding to one vs two copies of recombined *LoxP-STOP-LoxP-tomato* cassette. We found a slightly higher percentage of AT2s over Epcam in Exp. vs Ctrl lungs (37.7 vs 29.4%), indicating that the efficiency of labeling in AT2s in Exp. lung was not impaired compared to Ctrl lungs. The percentage of IAAPs over Epcam^Pos^ cells in Exp. vs Ctrl lungs (6.3% vs 7.6%, respectively) indicate similar labeling of these cells in Ctrl and Exp. conditions.

Altogether, these data indicate that the efficiency of labeling of IAAPs and AT2s in Ctrl and Exp. lungs is comparable at this earlier time point, suggesting that the increase in the IAAPs to AT2 ratio in Exp. lung at later time points is not due to a difference in the recombination efficiency of the *Rosa26* locus at earlier time points. We also compared the global efficiency of recombination at later time points in Exp. vs Ctrl by IF (without distinguishing between IAAPs and AT2s as this is not possible by IF using only Tomato) by quantifying the percentile of Tom^Pos^SftpcPos/SftpcPos at days 7 and 14. Our results indicate at day 7 similar proportion of Tom^Pos^SftpcPos/SftpcPos (d7: 77% ± 5.4 in Ctrl vs 70% ± 0.48 in Exp., n=4). Such observation was also made at day (d14: 84% ± 4.23 in Ctrl vs 82% ± 3.97 in Exp., n=4) (Figure 4b).

To investigate whether *Fgfr2b* was successfully deleted in both IAAPs and AT2s, RT-PCR was carried out to detect the wild type and mutant *Fgfr2b* transcripts (Figure 1c) [4]. The mutant *Fgfr2b* transcript (195 bp) was present in E-IAAPs and E-AT2s in *Fgfr2b-*cKO lungs, and as expected, was not detected in the corresponding cells (C-IAAPs and C-AT2s) from Ctrl lungs (Figure 1c). Sequencing of wild type and mutant cDNA bands that were cut and purified from agarose gel confirmed the deletion of exon 8 encoding the *Fgfr2b* isoform (Figure S3).

Next, qPCR was performed on FACS-isolated IAAPs and AT2s isolated from Ctrl and Exp. lungs. As previously described, we found that C-AT2s compared to C-IAAPs are enriched in *Fgfr2b*, *Etv5*, and the differentiation markers *Sftpc*, *Sftpb* and *S*ftpa1 (Figure 1d) [19]. However, in contrast to the Ctrl, *Fgfr2b* expression between E-IAAPs and E-AT2s is reduced. Moreover, *Etv5* expression is significantly downregulated in E-AT2s vs E-IAAPs and the expression levels of *Sftpc, Sftpb*, and *S*ftpa1 are not substantially different between E-IAAPs and E-AT2s (Figure 1e).

### ATAC-seq analysis and transcriptomic analyses reveal that *Fgfr2b* deletion leads to activation of IAAP cells

To carry out genome-wide profiling of the epigenomic landscape, an assay for transposase-accessible chromatin using sequencing (ATAC-seq) was performed on C-IAAP and E-IAAP subpopulations at day 7 on tamoxifen water (Figure S4). Interestingly, our data indicated a high signal background in E-IAAPs (data not shown), often seen in dying cells [26].

After correcting this elevated background to remove the contribution of dying cells, common and distinct peaks were identified for C-IAAPs and E-IAAPs (Figure S4a). Gene set enrichments based on regions of opened chromatin were carried out. Gene set enrichment (corrected P-value smaller than 0.2, top 50 sets) between C-IAAPs and E-IAAPs using Kobas PANTHER predicted that genes belonging to the inflammation mediated by chemokines and cytokine signaling pathway as well as genes controlling apoptosis signaling pathways were significantly upregulated in E-IAAPs compared to C-IAAPs (data not shown). Further analysis of the ATAC-seq data using the Reactome database indicated that the chromatin in loci of genes belonging to metabolism, metabolism of lipids and lipoprotein and immune genes was more open in E-IAAPs (Figure S4b).

We also explored using gene array carried out between C-AT2s, C-IAAPs and E-IAAPs captured at day 7 of tam water exposure, the status of the genes belonging to cell cycle, Fgfr2b transcriptomic signature previously identified [12] [13, 27] and the AT1/AT2 signature [28] (Figure 2). Our data confirm that C-IAAPs are quiescent cells compared to C-AT2s. However, we observe a drastic upregulation of cell cycle genes in E-IAAPs consistent with their activated status (Figure 2a). We also found that Fgfr2b signature at E12.5 [12] (Figure 2b), E14,5 [13] (Figure 2c) and E16.5 [27] (Figure 2d) is enriched in C-AT2s vs C-IAAPs. We also found these signatures to be upregulated in E-IAAPs, supporting the activation of Fgfr2b signaling in these cells (Figure 2b-d). Finally, we examined the status of the AT2 and AT1 signatures. As previously described, the C-AT2s are mature AT2s and express a high level of the AT2 signature compared to the undifferentiated C-IAAPs. By contrast, E-IAAPs display a significant enrichment in the AT2 signature, suggesting that these cells differentiate towards mature AT2s (Figure 2e).

**Fig. 2.**
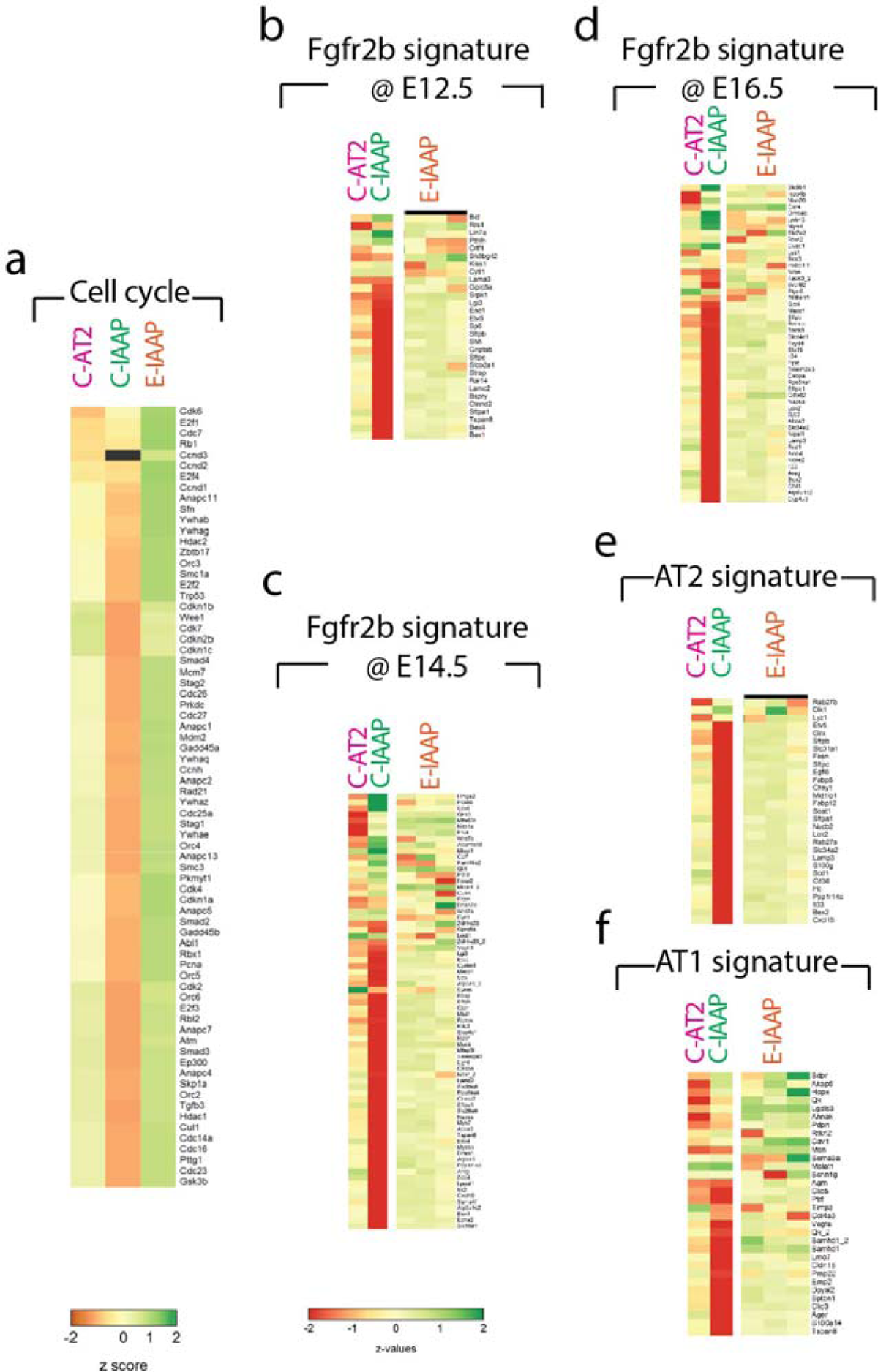
Gene arrays comparing C-AT2s, C-IAAPs and E-IAAPs. **a**) heatmap for the cell cycle genes indicating up-regulation of cell cycle genes in E-IAAP. **b**) heatmap for the Fgf10 signature at E12.5 indicating increased Fgf signaling in E-IAAPs. **c**) Heatmap for the AT2 signature supporting the increased commitment of the E-IAAPs towards the AT2 lineage

Interestingly, we found that the AT1 signature is also drastically increased in E-IAAPs vs C-IAAPs (Figure 2f). Comparison of the AT1 signature in C-IAAP vs C-AT2 reveals that only part of this signature is increased in C-IAAPs, but this increase includes bona fide AT1 markers such as *Hopx* and *Pdpn*. The emerging picture is therefore that E-IAAPs appear to display both an AT1 and an AT2 signature, suggesting that the lineage-labeled SftpcPos IAAP cells have the potential to engage into the AT1 lineage, a property that is well-accepted for mature AT2s.

Altogether, we conclude that the E-IAAPs analyzed on day 7 during tamoxifen water exposure are made of apoptotic and surviving cells. However, after background correction, a sub-population of E-IAAPs, highly active metabolically, likely proliferative, displaying increased Fgf signaling activation, enhanced AT2 and AT1 signatures and geared towards lipoprotein metabolism, which is associated with surfactant production, emerged.

### Fgfr2b inactivation in the AT2 lineage leads to the loss of Fgfr2b signaling in AT2s and activation of Fgfr2b signaling in IAAPs

The AT2s and IAAPs were compared between Exp. and Ctrl lungs using qPCR and immunofluorescence staining on cytospins of sorted cells (Figure 3a). qPCR analysis of AT2 demonstrated a significant decrease of *Fgfr2b* and *Etv5* expressions in Exp. vs Ctrl lungs, corroborating the loss of Fgfr2b signaling in these cells; however, no changes in the expression of *Sftpc, Sftpb*, and *S*ftpa1 was observed in these cells (Figure 3b). By contrast, in IAAPs, significant upregulation of *Fgfr2b*, *Etv5, Sftpc and Sftpb* was identified (Figure 3c). These changes in *Fgfr2b* and *Sftpc* mRNA levels were validated at the protein level by cytospin of isolated AT2s and IAAPs from Exp. and Ctrl lungs, followed by immunofluorescence staining (Figure 3d,e). These results support the loss of Fgfr2b signaling in AT2s and activation of Fgfr2b signaling in IAAPs in Exp. vs Ctrl lungs.

**Fig. 3.**
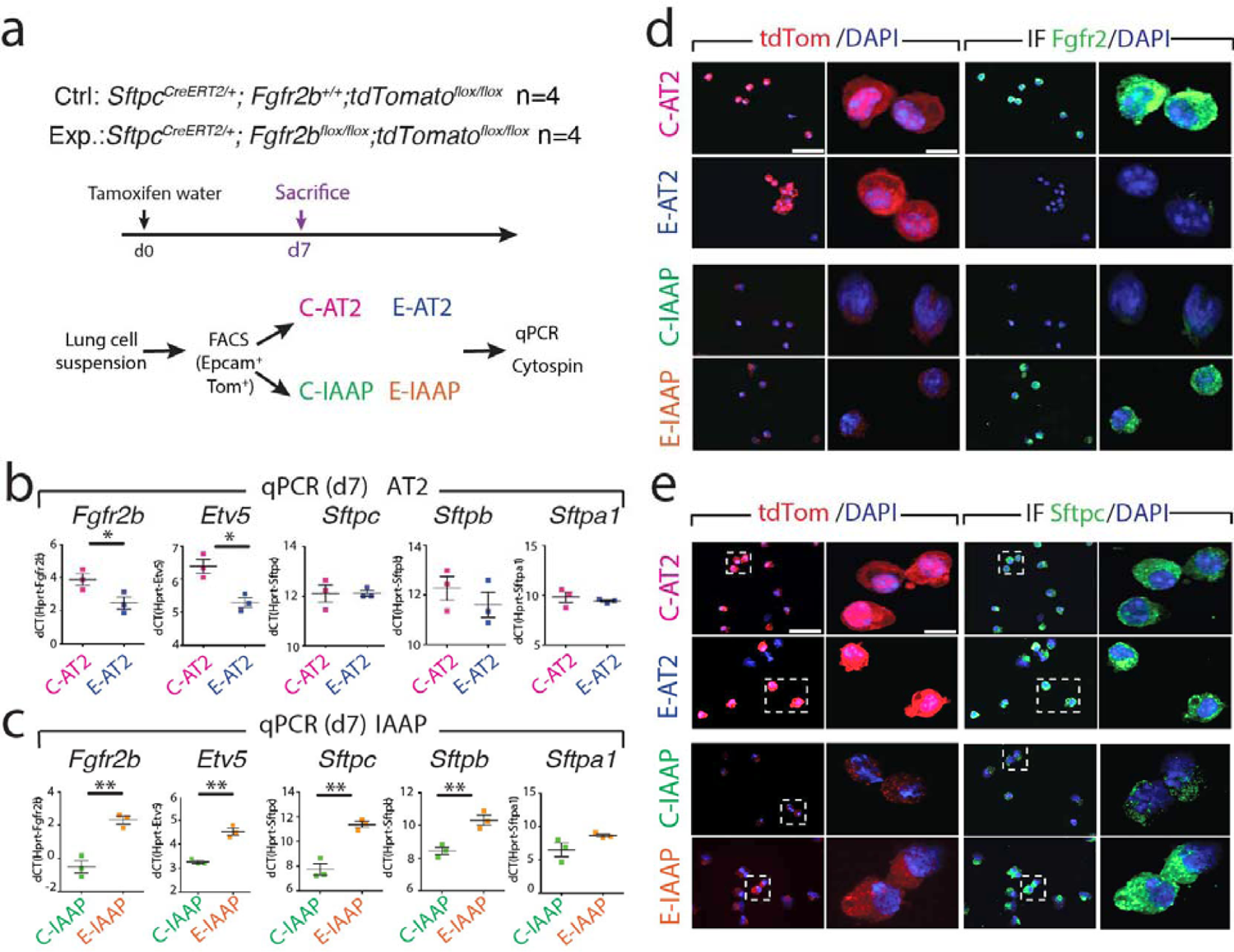
*Fgfr2b* inactivation in the AT2 lineage leads to the loss of Fgfr2b signaling in AT2s and activation of Fgfr2b signaling in IAAPs. **a)** Timeline of tamoxifen treatment of *S*ftpcCreERT2/^+^*; Fgfr2b^+/+^; tdTom^flox/flox^* and *S*ftpcCreERT2/^+^*; Fgfr2b ^flox/flox^; tdTom^flox/flox^* mice (n=4). **b)** qPCR gene expression analysis of FACS-based sorted C-AT2s and E-AT2s. **c)** qPCR gene expression analysis of FACS-based sorted C-IAAPs and E-IAAPs. **d)** Immunofluorescence staining against Fgfr2 on cytospins of C-AT2s, E-AT2s and C-IAAPs, E-IAAPs (Scale bar: 50μm). **e)** Sftpc Immunofluorescent staining on cytospins of C-AT2s, E-AT2s and C-IAAPs, E-IAAPs (n=4) (Scale bar: 50 SEM. Data are presented as mean values ± SEM. *p < 0.05, **p < 0.01, ***p < 0.001 μm).

### Genomic analysis in *Fgfr2b-*cKO reveals that the *Fgfr2b* locus is differentially impacted in mature AT2s and IAAPs

Given the surprising result that Fgfr2b signaling was activated in E-IAAPs despite the *Fgfr2b* deletion observed initially in these cells on day 7 after tamoxifen treatment (Figure 1e), the mice were treated for a longer time with tamoxifen to ensure that *Fgfr2b* deletion was complete. Next, the presence of the mutant and wild-type *Fgfr2b* transcripts was analyzed by RT-PCR on day 14 after tamoxifen treatment (continuous tamoxifen water treatment). Surprisingly, the results indicate that in E-IAAPs from *Fgfr2b-*cKO lungs, the mutated *Fgfr2b* transcript was barely detectable, while the mutated transcript was still detected in *Fgfr2b-*cKO E-AT2s at both time points (Figure 4a,b).

**Fig. 4.**
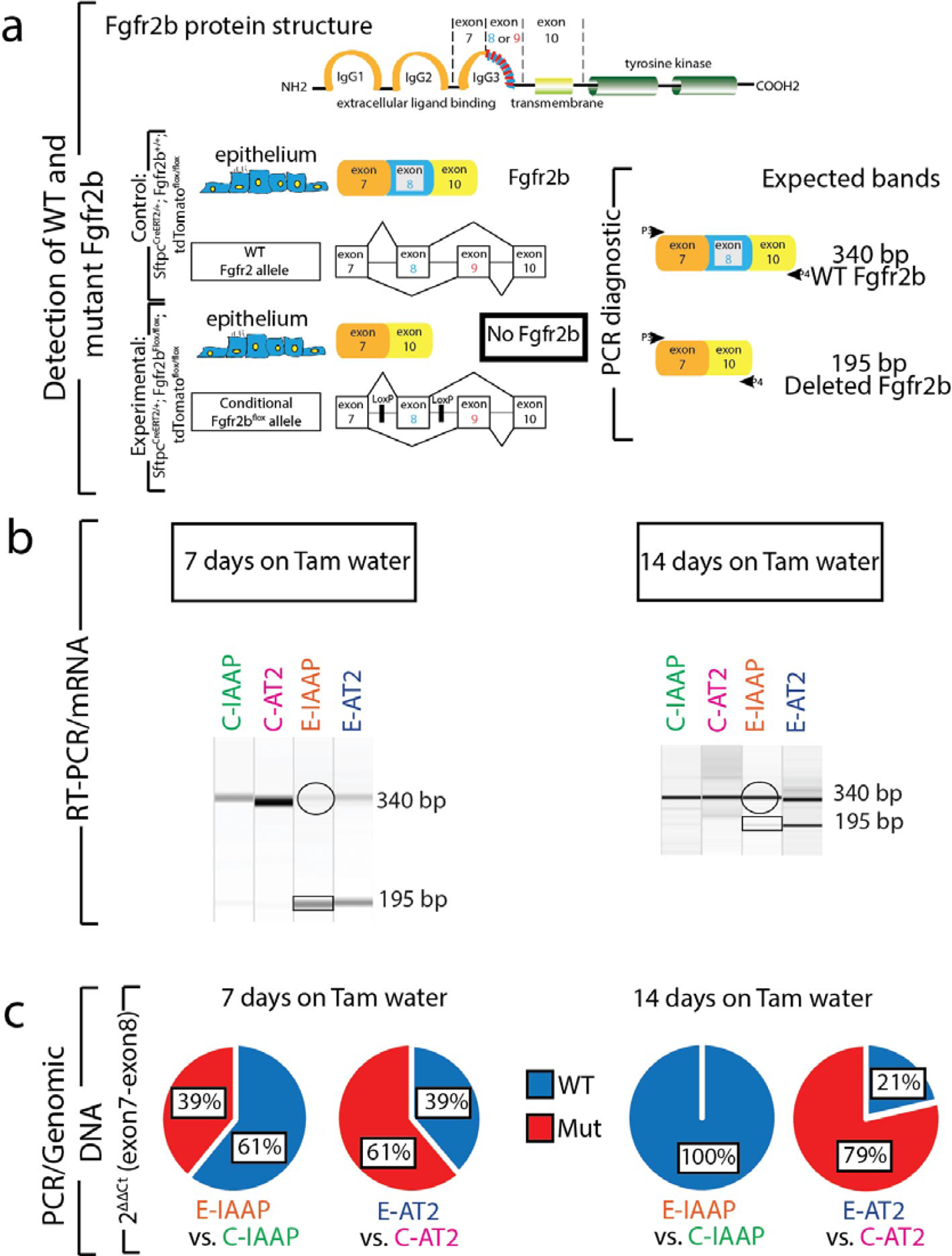
Continuous deletion of the *Fgfr2b* allele in AT2 cells and amplification of IAAP cells. **a)** Schematic of Fgfr2b protein structure, coding mRNA and DNA. Wild type *Fgfr2b* transcript consists of exon 7, exon 8 and exon 10, which is detected by the band size of 340bp, and mutant *Fgfr2b* form (exon 8 deleted) is detectable by the band size of 195bp. **b)** RT-PCR for detecting WT and *Fgfr2b* mutant transcripts in FACS-based sorted C-IAAPs, C-AT2s, E-IAAPs and E-AT2s on day 7 and day 14. **c)** Pie charts represent qPCR data for deleted exon 8 (mutated Fgfr2b locus) vs exon 7 (reflecting the intact *Fgfr2* locus) to detect the relative extent of the mutated and wild type *Fgfr2b* locus in E-IAAPs vs C-IAAPs and E-AT2s vs C-AT2s at two time points

To quantify the mutated and wild type *Fgfr2b* on day 7 and day 14, qPCR for the detection of exon 8 (the deleted exon) vs exon 7 (reflecting the intact *Fgfr2b* locus) was performed on the genomic DNA of AT2s and IAAPs from *Fgfr2b-*cKO and Ctrl lungs. The results show that on day 7, the relative presence of the mutated and wild type *Fgfr2b* in E-IAAPs was 39% and 61%, respectively. However, on day 14, wild type *Fgfr2b* increased to 100% while the mutated Fgfr2b was no longer detected, indicating that at this time point E-IAAPs contain mostly the wild type *Fgfr2b* (Figure 4c). In E-AT2s, by contrast, there was an increase in the percentage of mutated *Fgfr2b* from day 7 to day 14 (from 61% to 79%), indicating that continuous deletion of the *Fgfr2b* allele in AT2 cells occurs. This result supports the amplification of E-IAAPs containing wild type *Fgfr2b* and the continuous deletion of *Fgfr2b* in E-AT2s. However, the molecular mechanisms involved in the expansion of the E-IAAPs with the wild type *Fgfr2b* allele are still unclear. One possibility is that the previously described low level of *Sftpc* (which should translate into a lower level of Cre recombinase) associated with the closed chromatin configuration in IAAPs renders difficult the efficient recombination of the exon 8 of the *Fgfr2b* locus.

### Reduction of tdTomato^Pos^ cells along with enhanced apoptosis and proliferation in Exp. *Fgfr2b-*cKO

FACS analysis of the percentage of tdTomato^Pos^ over Epcam^Pos^ in Ctrl and *Fgfr2b-*cKO indicates the reduction of tdTomato labeled cells following *Fgfr2b* deletion on day 7 and day 14 (Figure 5a,b). Quantification of tdTom^Pos^ cells over total (DAPI^Pos^) cells on sections supports this result (Figure 5c).

**Fig. 5.**
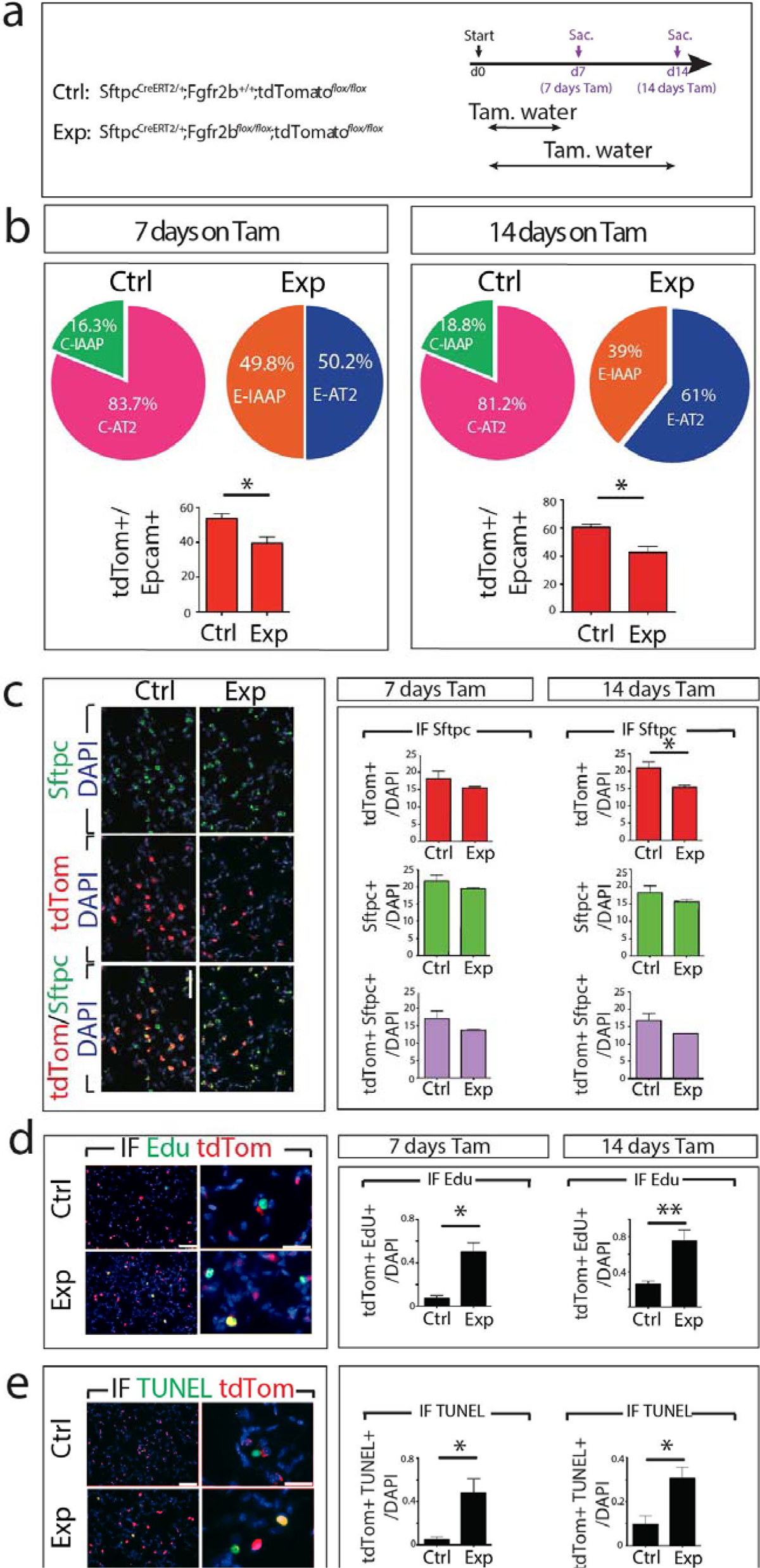
Reduction of tdTomato^Pos^ cells along with enhanced apoptosis and proliferation in *Fgfr2b-*cKO. **a)** Tamoxifen treatment timeline of *S*ftpcCreERT2/^+^*; Fgfr2b^+/+^; tdTomato^flox/flox^* and *S*ftpcCreERT2/^+^*; Fgfr2b^flox/flox^; tdTomato^flox/flox^* mice. **b)** Flow cytometry analysis of the percentage of tdTomato^Pos^ cells in Ctrl and Exp. on day 7 and day 14. Note the expansion of the E-IAAPs as well as the decrease in tdTom+/Epcam+ in Exp. lungs. **c)** Representative Sftpc immunofluorescence staining (Scale bar: 50 m). Quantification of tdTomato+, Sftpc+ single positive and tdTomato+ Sftpc+ double-positive cells at day 7 and day 14 (n=4). **d)** Representative EdU staining (Scale bar: 50μm) and quantification of tdTomato+ Edu+ cells at day 7 and day 14 (n=4). **e)** Representative TUNEL staining and quantification of tdTomato+TUNEL+ cells on day 7 and day 14. Data are presented as mean values ± SEM. *p < 0.05, **p < 0.01, ***p < 0.001

In addition, Sftpc IF staining of Ctrl and Exp. lungs was performed and quantified (Fig. 5c). The results indicate a trend towards a decrease of SftpcPos tdTom^Pos^ over total cells in *Fgfr2b-*cKO compared to Ctrl on days 7 and 14.

We also investigated proliferation and cell death of tdTom^Pos^ cells by immunofluorescence staining on lung sections on days 7 and 14 (Figure 5d,e). In this context and as previously reported [19], Tomato fluorescence on sections does not distinguish between IAAPs and AT2s. On days 7 and 14, a significant increase in proliferation (Figure 5d) and apoptosis (Figure 5e) in tdTom^Pos^ cells was observed, suggesting that lineage-labeled subpopulations undergo apoptosis and proliferation simultaneously in Exp. lungs. These results, combined with the expansion of the IAAPs and the loss of AT2s in Exp. lungs, indicate that upon *Fgfr2b* deletion the IAAPs proliferate while the AT2s die.

### The lung structure remains Intact following *Fgfr2b* deletion in the AT2 lineage

To investigate whether there is a change in the lung structure after *Fgfr2b* deletion, lung morphometry analysis was performed on days 7 and 14 after tamoxifen treatment. Our results demonstrate no changes in alveolar space, septal wall thickness and MLI in *Fgfr2b-*cKO compared to Ctrl (Figure 6a-e). These results suggest that the lack of abnormal lung phenotype is linked to a continuous compensatory mechanism that replenishes the mature AT2 pool. Therefore, we hypothesized that IAAPs, as immature AT2 cells, are the cells that proliferate and differentiate to mature AT2 cells.

**Fig. 6.**
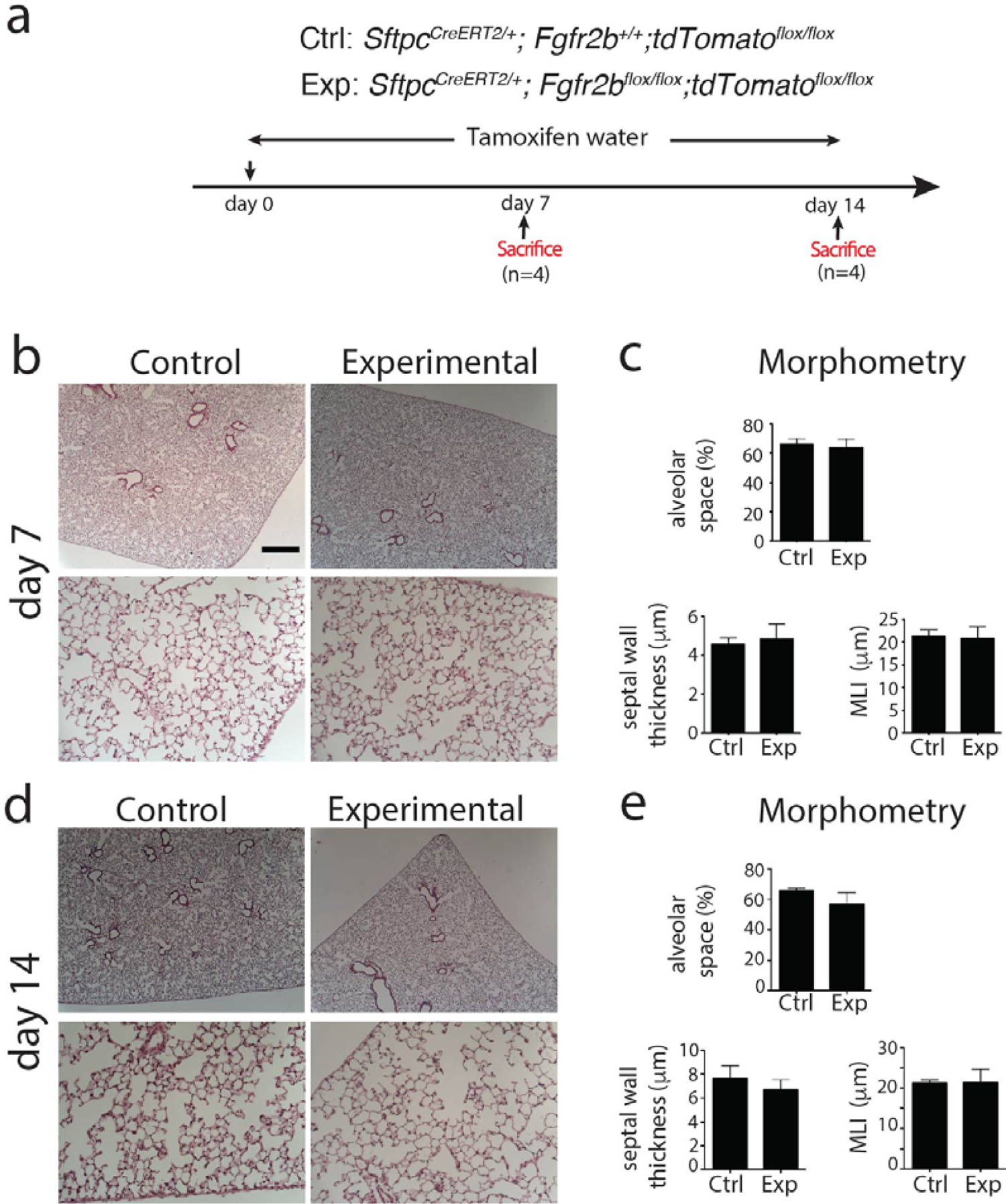
Lung structure remains Intact following *Fgfr2b* deletion. **a)** Timeline of tamoxifen treatment of *S*ftpcCreERT2/^+^*; Fgfr2b^+/+^; tdTomato^flox/flox^* and *S*ftpcCreERT2/^+^*; Fgfr2b^flox/flox^; tdTomato^flox/flox^* mice. **b)** Hematoxylin and eosin staining of the Ctrl and the Exp. lungs at day 7 (scale bar 200 and 50 μm) **c)** Morphometry analysis (alveolar space, septal wall thickness, and MLI) of the Ctrl and the Exp. lungs at day 7 (n=4). **d)** Hematoxylin and eosin staining of the Ctrl and the Exp. lungs at day 14 (scale bar 200 and 50 μm). **e)** Morphometry analysis (alveolar space, septal wall thickness, and MLI) of Ctrl and Exp. lungs at day 14 (n=4). Data are presented as mean values ± SEM. p < 0.05, p < 0.01, p < 0.001

### Deletion of *Fgfr2b* in the AT2 lineage leads to loss of self-renewal capability in mature AT2s and a gain of alveolosphere formation potential in IAAPs

To compare the proliferative capacity of IAAPs and AT2s in Ctrl and *Fgfr2b-*cKO lungs, FACS-based sorted cells were co-cultured with Cd31^Neg^Cd45^Neg^Epcam^Neg^Sca1^Pos^ resident lung mesenchymal cells according to a previously published protocol (Figure 7a). AT2s from Ctrl lungs behaved as *bona fide* AT2 cells as they formed alveolospheres with the expected colony-forming efficiency (Figure 7b,c). By contrast, AT2s from *Fgfr2b-*cKO lungs demonstrated a significant decrease in alveolosphere forming capabilities compared to the corresponding Ctrl, suggesting the loss of proliferative capabilities upon *Fgfr2b* deletion (0.22% ± 0.13 vs 1.20% ± 0.36, n=3) (Figure 7c). As previously described, IAAPs from Ctrl lungs displayed weak organoid forming capabilities, which is in line with their quiescent status [19]. Interestingly, IAAPs from *Fgfr2b-*cKO lungs showed a significant increase in alveolosphere formation, which is consistent with their transition towards the AT2 status (0.02% ± 0.01 vs 0.15% ± 0.05, n=3) (Figure 7d,e). Supporting this conclusion, we observed differential viability of FACS-isolated AT2s and IAAPs from Ctrl and *Fgfr2b-*cKO lungs. AT2s displayed decreased viability in Exp. vs Ctrl lungs (18,27% ± 1,64% vs 72,33% ± 5,62%, n=3). By contrast, a sharp increase in viability was observed for IAAPs in Exp. vs Ctrl lungs (72% ± 4% vs 10,17% ± 0,98%, n=3) (Figure S5). These results suggest that IAAPs in *Fgfr2b-*cKO lungs display progenitor behavior characteristics similar to mature AT2s in the Ctrl lungs. Indeed, such progenitor-like behavior was previously suggested in vitro. Using precision-cut lung slides from [*S*ftpcCreERT2/^+^*; Fgfr2b^+/+^; tdTom^flox/flox^*] mice cultured in vitro, we demonstrated that mature AT2s are lost while IAAPs are expanded. In vivo, we also showed that the IAAPs are expanding following pneumonectomy [19].

**Fig. 7.**
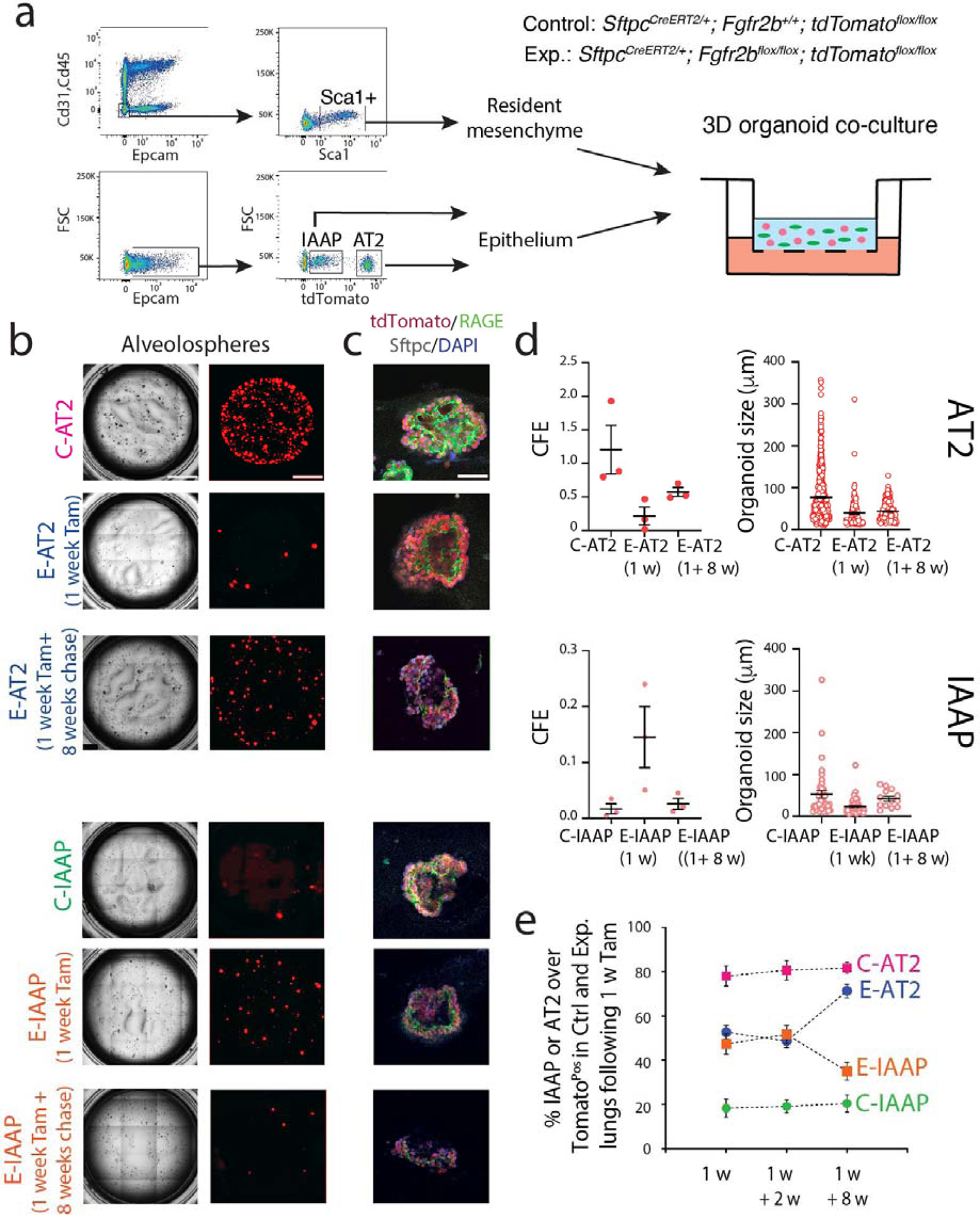
Deletion of *Fgfr2b* in the AT2 lineage leads to loss of self-renewal capability in AT2 and a gain of alveolosphere formation potential in IAAPs. **a)** Representative flow cytometry shows the gating strategy of Cd31^Neg^Cd45^Neg^Epcam^Neg^population and a further selection of Sca1+ resident mesenchymal cells from C57BL/6 lungs (upper plot), as well as the selection of IAAPs and AT2s from Epcam^Pos^ population from *S*ftpcCreERT2/^+^*; Fgfr2b^+/+^; tdTomato^flox/flox^* (lower plot). Resident mesenchymal cells were co-cultured with IAAPs and AT2s separately (n=3). **b)** Representative alveolospheres from AT2s and IAAPs from Ctrl and Exp. mice (n=3), (Scale bar: 100μm) **c)** Representative Sftpc and RAGE immunofluorescence staining of alveolospheres after 14 days in culture, (Scale bar: 50μm). **d)** Quantification of alveolospheres size and Colony-forming unit (CFU) in AT2s and IAAPs from Ctrl and Exp. mice (n=3). **e)** Percentile of AT2s and IAAPs in Ctrl and Exp lungs at different time points following 7 days tamoxifen treatment.

### E-AT2s regain their alveolosphere formation capabilities following a long chasing period after Tamoxifen exposure

We also tested the capacity of the E-AT2 and the E-IAAPs to give rise to alveolospheres in Exp. mice exposed to Tam water for one week followed by a chase period of 8 weeks. In these conditions, a partial but significant rescue of the capacity of the E-AT2 was observed compared to the E-AT2 arising from animals exposed to Tam water for one week. On the other hand, the E-IAAPs lost their proliferative activity after such a long chase period compared to the E-IAAPs isolated from one-week tamoxifen treatment. The ratio of IAAPs or AT2s over the total Tom^Pos^ cells after one-week tamoxifen water, followed by a two-week or eight-week chase period, is represented in Figure 7e. Our results indicate that for the one-week and one-week plus two-week chase period, the percentile of E-AT2 and E-IAAPs is roughly equivalent and around 50%, down from 80% for the C-AT2s, up from 18% for the C-IAAPs. However, for the one-week tamoxifen plus eight-week chase period, these percentiles have almost returned to normal. The complete return to normality after an eight-week chase is likely hampered by the previously reported leakiness of the *S*ftpcCreERT2 driver, which in experimental mice continuously deletes *Fgfr2b* in AT2s arising either from de novo targeted AT2s or from the IAAPs which have differentiated into AT2s (see also Figure 9). Interestingly, such a dynamic mechanism was also observed in the context of bleomycin injury in mice, where at day 14 following bleomycin administration (at the peak of fibrosis), the AT2s decrease while the IAAPs simultaneously increase. On day 28, when the fibrosis resolution process has taken place, the percentile of IAAPs and AT2s have almost normalized (Zhang and Bellusci, data not shown_Please see Supplementary Figure for reviewers only).

### Transition of IAAPs towards AT2s in response to *Fgfr2b* deletion

The average expression of tdTomato intensity in the IAAP cells, obtained by flow cytometry in Ctrl and *Fgfr2b-*cKO lungs was quantified (Figure 8a). Our results indicate an expansion of the IAAPs in *Fgfr2b-*cKO vs Ctrl lungs towards higher tdTomato intensity. Furthermore, quantifying the mean fluorescence intensity in IAAPs in *Fgfr2b-*cKO vs Ctrl lungs confirmed this increase (Figure 8b).

**Figure 8:**
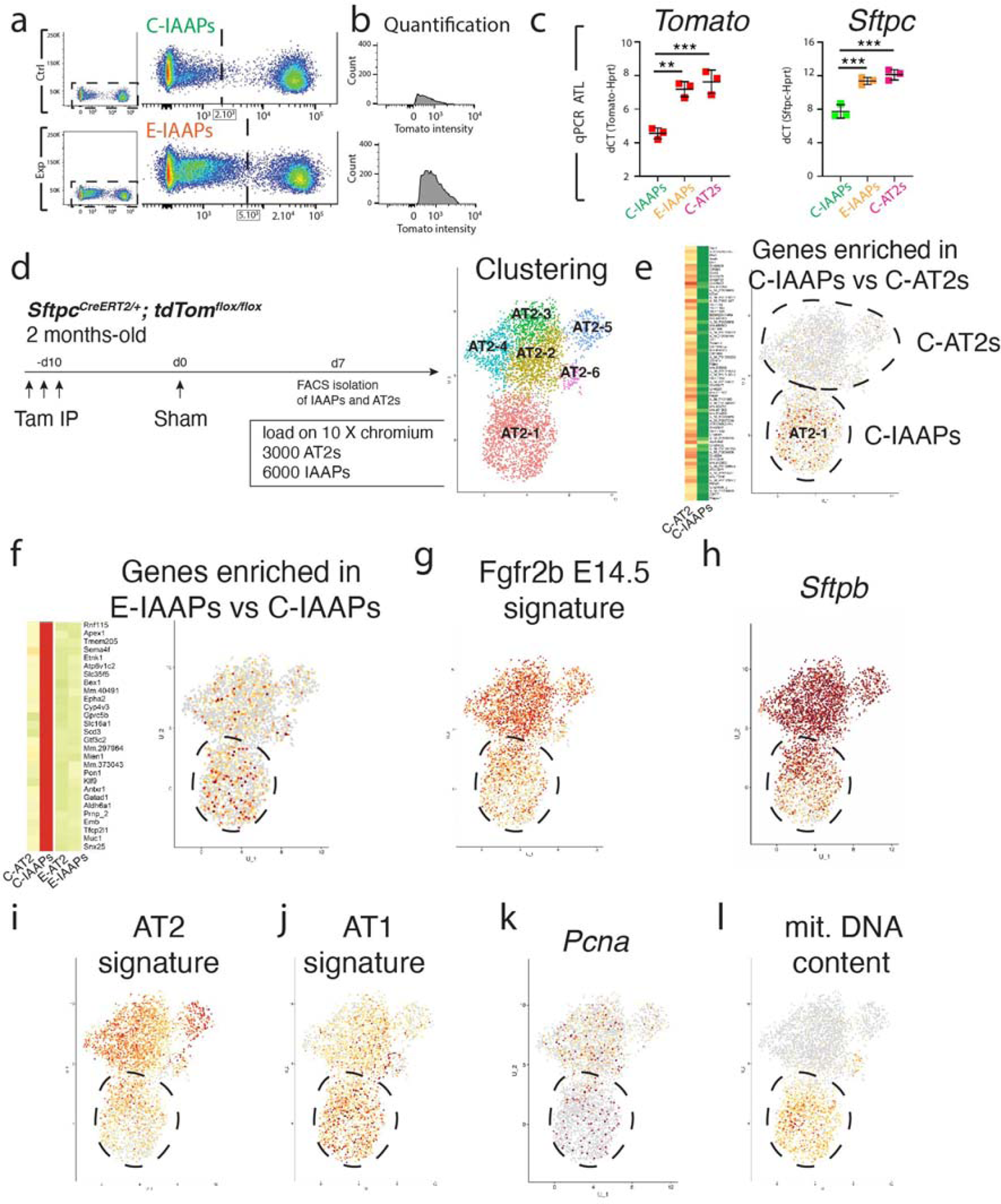
Transition of IAAPs towards AT2 in response to *Fgfr2b* deletion. **a)** Representative flow cytometry analysis of tdTomato shows the expansion of IAAPs towards higher tdTomato intensity in E-IAAPs compared to C-IAAPs. **b)** tdTomato intensity quantification of IAAPs in Ctrl and *Fgfr2b-*cKO lungs. **c)** qPCR analysis of *tdTomato* expression on FACS-based sorted IAAPs. **d**) scRNA-seq on FACS-isolated IAAPs and AT2s 7 days following Sham surgery. UMAP clustering indicates 6 main clusters. **e**) Expression of genes enriched in C-IAAPs vs. C-AT2s identifies the cluster AT2-1 as the IAAPs. **f**) Expression of genes enriched in E-IAAPs vs, C-IAAPS indicate that AT2-1/IAAPs subcluster contains activate IAAPs. **g**) Expression of genes representing the Fgfr2b E14.5 signature is enriched in AT2s. **h**) Expression of *Sftpb* is enriched in AT2s. **i**) Expression of genes representing AT2 signature is enriched in AT2s. **j**) Expression of genes representing AT1 signature is enriched in IAAPs. **k**) Expression of *Pcna* is present in both AT2s and IAAPs. **l**) Expression of mitochondrial DNA genes in IAAPs. Data are presented as mean values ± SEM. ∗p < 0.05, ∗∗p < 0.01, ∗∗∗p < 0.001.

Next, the level of expression of *Tomato* mRNA in FACS-isolated IAAPs in *Fgfr2b-* cKO vs C-IAAPs and C-AT2s in Ctrl lungs was quantified and compared by qPCR. We found a substantial upregulation of *Tomato* expression upon *Fgfr2b* deletion (Figure 8c), suggesting that in the *Fgfr2b-*cKO lungs, the IAAPs are transitioning towards an AT2 status. Interestingly, ATAC-seq analysis indicated more open chromatin, in E-IAAPs vs C-IAAPs, in the *Rosa26* locus, containing the *tdTomato* gene (data not shown). These results are also in line with the qPCR analysis of the AT2 cell differentiation *Sftpc* indicating increased expression in E-IAAPs vs C-IAAPs and a level of expression close to the one observed in C-AT2s (Figure 8c)

### ScRNA-seq analysis of the AT2 lineage demonstrates that IAAPs and mature AT2s exist as two independent but related clusters

Next, we used scRNA-seq to expand the profiling of IAAPs and AT2s beyond the bulk population analysis done previously. In particular, scRNA-seq allows defining better the level of heterogeneity present in given populations (Fig. 8d-l). As we previously described that IAAPs get activated and proliferate upon pneumonectomy, we isolated the IAAPs and AT2s on day 7 after sham or PNX. The results presented below focus only on the sham, which we considered a surrogate for Ctrl lungs.

First, we used flow cytometry to sort separately IAAPs and AT2s cells from sham lungs (obtained from pooling these cells from 3 mice). As the C-IAAPs were described as more fragile than the C-AT2s following flow cytometry, we loaded on the 10X chromium chip a total of 9000 cells made of 3000 C-AT2s and 6000 C-IAAPs. Fine clustering allowed us to distinguish 6 clusters (AT2-1 to AT2-6) (Figure 8d). Then, the lineage-labeled cluster(s) which corresponded to the IAAPs was identified by interrogating the transcriptomic signature (arising from bulk RNAseq) obtained by comparing C-IAAPs and C-AT2s [19]. Our results indicated that the AT2-1 cluster displayed a high level of IAAPs signature compared to the other clusters (Figure 8e).

We also monitored the presence of a transcriptomic signature enriched in E-IAAPs vs C-IAAPs (Figure 8f). This signature is also normally massively decreased in C-IAAPs vs C-AT2s. However, the cells in the AT2-1/IAAPs cluster in our scRNA-seq displayed a higher level of this signature than the AT2s suggesting that the IAAPs arising from sham lungs display similar transcriptomic profiles as the E-IAAPs and are therefore activated (Figure 8f).

Next, we examined the Fgfr2b transcriptomic signature at E14.5 and found it to be present in the AT2-1/IAAPs cluster, albeit at a lower level compared to the AT2s cluster (Figure 8g). Consistent with previous results, we also found that *Sftpb* expression was decreased in IAAPs vs AT2s (Figure 8h). A similar observation was done with the AT2 transcriptomic signature (Figure 8i). Confirming the bulk population analysis (Figure 2f), we found that the AT1 signature was also significantly increased in IAAPs vs AT2s. (Figure 8j). We also found that the IAAPs contained cells expressing *Pcna* and a higher level of mitochondrial content (Figure 8k,l), thereby supporting the previous observation that IAAPs were proliferative and metabolically active.

Interestingly, we did not find a large number of *Pd-l1* (*Cd274*) expressing cells in our data set, suggesting that either these cells did not survive in our experimental conditions (even though roughly 50% of the IAAPs should be expressing Pd-l1 both at the protein and mRNA level). The alternative possibility is that these cells lost *Pd-l1* mRNA expression during scRNA-seq. In our experimental conditions, the time separating the isolation of the lungs from the loading of the cells on the 10X Genomic chip, which is known to influence gene expression, was around 5 hours.

In summary, we have demonstrated that the IAAPs and AT2s represent two transcriptionally stable and distinct populations of SftpcPos cells. Fine clustering indicated heterogeneity in AT2s (with five subclusters). However, such heterogeneity was not detected in the IAAPs. It is still unclear if such a result is because initial IAAPs subpopulations are excluded from the analysis process due to their fragility or if the IAAPs change their transcriptomic profile over time during the scRNA-seq process. More work is needed to clarify this critical aspect.

## DISCUSSION

The model for our study is presented in Figure 9. Using the *S*ftpcCreERT2/^+^*; tdTomato^flox/+^* mice, we have previously described the existence of two distinct subpopulations of lineage-traced SftpcPos cells based on the level of Tomato expression. The AT2-Tom^High^ represent the mature AT2 cells. On the other hand, the AT2-Tom^Low^ displayed characteristics of immature AT2 cells that could proliferate and differentiate towards mature AT2 cells in the context of pneumonectomy. These cells were proposed to represent a novel progenitor population for mature AT2 cells [19]. Due to these characteristics, we are calling them “injury-activated alveolar progenitors” or IAAPs. In the context of *Fgfr2b* deletion, both the AT2s and IAAPs undergo apoptosis. Our ATAC-seq data supports these results showing a much higher background noise in E-IAAPs on day 7 compared to the corresponding Ctrl. Such high background noise has been associated with apoptosis [26]. In addition, IF data showed an increase in the tdTom^Pos^ TUNEL^Pos^ cells. Unfortunately, IF for Tomato does not distinguish between AT2s and IAAPs on sections [19]. As E-AT2 cells have lost their proliferative capabilities in the context of the alveolosphere assay, AT2s are likely the most affected cells by the loss of *Fgfr2b*. This conclusion is supported by the high expression level of the mutated transcript in E-AT2s on day 7 of tamoxifen treatment. In addition, while in the AT2 pool, this apoptotic phenotype is fully penetrant. In the IAAP pool, we observed the emergence of lineage-labelled IAAP cells that did not display *Fgfr2b* deletion. This result suggests that in these IAAP cells, Cre would act preferentially in the *Rosa26* locus to activate Tomato expression but would not operate efficiently on the *Fgfr2b* locus to delete the exon 8. These cells would therefore represent transient amplifying cells with progenitor-like properties.

**Figure 9:**
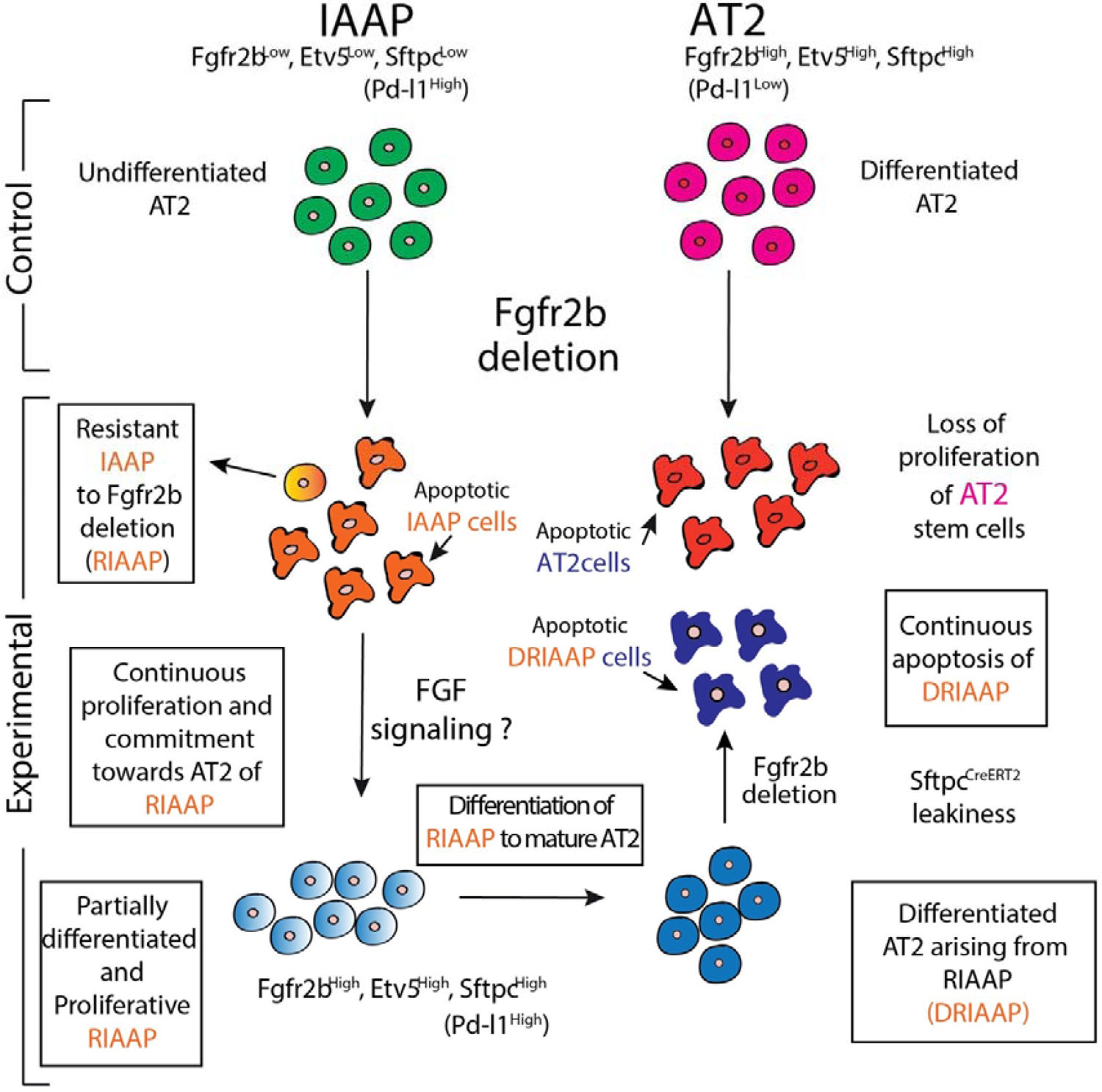
Schematic representation of characteristics and behavior of IAAP and AT2 cells in Ctrl and *Fgfr2b-*cKO lungs. The AT2-Tom^High^ are the mature AT2s while the AT2-Tom^Low^ (IAAPs) correspond to immature AT2s. In the context of *Fgfr2b* deletion, both AT2s and IAAPs undergo apoptosis. However, while in the AT2 pool, this apoptotic phenotype is fully penetrant, in the IAAP pool, we observed the emergence of resistant IAAPs to *Fgfr2b* deletion (RIAAPs). These results also suggest that the IAAP pool is itself heterogeneous. The difference between RIAAPs and IAAPs and the mechanisms involved in the emergence of this resistance in RIAAPs will require further investigation. We also propose that the RIAAPs proliferate and get progressively committed towards mature AT2s. We suggest that Fgf signaling in these cells is likely driving this proliferation and differentiation process. Differentiated AT2 arising from RIAAPs (DRIAAPs) are then, due to previously described leakiness of the *S*ftpcCreERT2 driver ^19^, undergoing *Fgfr2b* deletion creating a constant cycle of proliferative and apoptotic alveolar epithelial cells allowing to maintain AT2 homeostasis. The long-term consequences of this new equilibrium are still unclear. In addition, how different are the DRIAAPs from bona fide AT2s is still unknown.

Interestingly, a disconnect between Tomato expression (serving as quality control for Cre activity) and Cre-induced Diphtheria Toxin (DTA) activity in AT2 cells has been previously reported [20]. The authors used *S*ftpcCreERT/^+^*, R26^LoxP-STOP-LoxP-Tomato^; R26^LoxP-GFP-STOP-LoxP-DTA^* to label the AT2 cells and induce in the same time the lethal expression of DTA in these cells after a single dose of tamoxifen. It was observed that lineage-labeled AT2 cells (which generally should have died due to DTA expression) proliferated clonally following AT2 killing. These data served as a base to demonstrate that AT2 are stem cells. To explain their results, the authors proposed that “by chance, Tamoxifen-induced recombination occurred only at the *Rosa^LoxP-STOP-LoxP-Tomato^* locus in a proportion of AEC2s (AT2s), thereby lineage labeling, but not killing these AEC2s (AT2s)” [20]. A puzzling possibility in this experiment is that the lineage-labeled AT2 cells which proliferated clonally following AT2 killing arise from the lineage-labelled IAAPs.

The interpretation of these results is consistent with our published observation that in the context of precision-cut lung slices from *S*ftpcCreERT2/^+^*; tdTomato^flox/flox^* lungs cultured in vitro, AT2s are massively killed, leaving intact the IAAPs, which then expand to become mature AT2s [19]. In the context of *Fgfr2b* deletion, rather than random recombination of one allele vs the other upon tamoxifen administration, an alternative scenario is that a subset of lineage-labelled IAAPs are or become resistant to *Fgfr2b* deletion, allowing them to survive. We call these cells <u>R</u>esistant IAAP cells to *Fgfr2b* deletion (or RIAAP cells). The mechanisms involved in the resistance in RIAAP cells will require further investigation. Novel mechanisms are likely at play as this observation is not compatible with a simple difference in chromatin opening, restricting, for example, the accessibility of the *Fgfr2b* locus. A closed chromatin configuration for the *Fgfr2* locus would expect to hamper both the recombination of the *exon 8* and the expression of *Fgfr2b* itself. As Fgfr2b expression is, on the contrary, increased in RIAAPs, it is clear that this primary epigenetic mechanism is not sufficient to explain our results. These results also suggest that the IAAP pool is itself heterogeneous, and the difference between surviving RIAAPs and dying IAAPs will need further clarification. We also propose that the RIAAPs proliferate and get progressively committed towards mature AT2s. Based on the increased expression of *Fgfr2b* and *Etv5* and the AT2 differentiation markers, we propose that Fgf signaling in these cells is likely driving the proliferation and differentiation process.

Further experiments will have to be carried out to identify the Fgf ligand, likely Fgf7 or Fgf10, driving these processes. These <u>d</u>ifferentiated AT2s arising from RIAAPs (called DRIAAPs) are then, due to the previously described leakiness of the *S*ftpcCreERT2 driver [19], undergoing *Fgfr2b* deletion creating a constant cycle of proliferative and apoptotic alveolar epithelial cells allowing to maintain AT2 homeostasis. It is also essential to consider that non-lineage labeled AT2s are still present in the Exp. lung. In our conditions, our labeling efficiency of AT2 cells is around 77% [19]. Therefore, it is possible that in the E-AT2 pool, there is a mixture of cells arising from E-IAAPs and cells de novo arising from non-lineage labeled AT2s, which undergo Cre-based recombination in a tamoxifen independent manner.

Indeed, the leakiness of the Cre in the SftpcCreERT2 line used for our study is relatively high and gives rise to around 5% of Tom^Pos^ cells/total cells labeled in mice exposed to water compared to 25% in the context of tamoxifen water [19]. The long-term consequences of this new equilibrium are still unclear. In addition, how different are the DRIAAPs from bona fide AT2s is still unknown. The overall effect of such a process triggered by *Fgfr2b* deletion in AT2s and IAAPs is a zero-sum game in terms of the appearance of a deleterious, emphysematous-like phenotype.

It was previously reported that mutant mice displaying specific deletion of Fgfr2 in AT2 cells were less prone to repair after injury, displayed enhanced mortality, and had reduced AT2 cells overall [29]. During homeostasis, *Fgfr2* deletion resulted in increased airspace and collagen deposition, as well as a reduced number of AT2 cells supporting our result that Fgfr2 is essential for AT2 maintenance. Earlier work investigating the consequences of the loss of *Etv5* in AT2 cells during homeostasis and repair after bleomycin-induced lung injury showed that Etv5 is required to maintain AT2 cells [30]. Upon *Etv5* deletion, AT2s transdifferentiated to AT1s. Furthermore, the repair process of the epithelium after lung injury was impaired, resulting in fewer AT2 cells altogether. As Etv5 is regulated by Fgfr2b signalling [31], it was suggested that Etv5 in AT2 cells is controlled by Ras-mediated ERK signalling [30].

A recent paper reported also the inactivation of *Fgfr2* in AT2s using the *S*ftpcCreERT2 driver line [14]. Their study concluded that Fgfr2 signaling is dispensable during homeostasis in the adult while it prevents the differentiation of AT2s towards the AT1 lineage during alveologenesis. In our conditions, loss of Fgfr2b signaling in AT2s leads to a significant decrease in their proliferative capacity using the alveolosphere assay. This result was not observed by Liberti et al. [14]. A methodological difference that could explain these results is that a different tamoxifen regimen was used. While we primarily studied the impact of *Fgfr2b* deletion on day 7 from the start of tamoxifen delivery via water, Liberti et al. treated the Exp. adult mice by oral gavage with tamoxifen for three consecutive days followed by two weeks washout period. We have also analyzed the lungs after a two-week (Figure S6) or eight-week (Figure S7) chase period. We observed increased proliferation and apoptosis in Tom^Pos^ cells at both time points, indicating the establishment of another homeostatic equilibrium. RT-PCR after one week of tamoxifen water followed with a two-week chase period shows that the WT transcript is dominating in E-AT2s and E-IAAPs (Figure S6a).

Interestingly, we also observe increased proliferative capabilities of the E-AT2 cells in the context of one-week tamoxifen followed by an eight-week chase period compared to the one-week tamoxifen water treatment (Figure 7). However, their capacity to proliferate is nonetheless decreased compared to C-AT2s. The difference between our conditions and the ones from Liberti et al. could be due to different leakiness levels of the *S*ftpcCreERT2 driver used.

Interestingly, Liberti et al. also reported, using IF, an increase in Edu^Pos^ lineage-traced cells in the Exp. vs Ctrl lungs without providing a clear explanation for this controversial result. Fgfr2b signaling is known to control proliferation or/and survival positively. However, to our knowledge, it does not inhibit proliferation *per se*. The interpretation for this puzzling result is now clear if we consider the proliferative lineage-traced E-IAAPs and the new homeostatic equilibrium present in the Exp. lungs.

We have also reported that C-IAAPs also express Pd-l1 [19]. We found similar results for E-IAAPs (Figure S8). In the context of cancer, PD-L1 expressed by some human cancer cells binds to PD1, a checkpoint protein expressed by T cells to prevent the immune cells from attacking them, allowing the cancer cells to escape the immune aggression. These cells usually display enhanced self-renewal capabilities and are considered cancer stem cells [21, 32–34]. A similar concept is emerging in the context of the IAAPs with their capacity to escape the harmful consequences of *Fgfr2b* inactivation. While these escaping properties may be beneficial in lung injury, future research should also focus on examining their role in the context of cancer. Designing dual-labeling systems such as the Dre/Rox and Cre/LoxP system [35] under the control of *Sftpc* and *Pd-l1* promoter appears to be a promising strategy to specifically target the IAAPs and examine their precise contribution to the AT2 lineage in the context of repair after injury, regeneration or even cancer.

In conclusion, we have identified IAAPs as a potentially novel population of AT2 progenitors necessary for alveolar repair after massive injury to mature AT2s. Understanding how IAAPs get activated to proliferate and differentiate into mature AT2s will be critical to designing efficient strategies to treat debilitating lung diseases.

## Supporting information

supplementary data for Ahmadvand et al

## DECLARATIONS

### Ethics approval

All animal studies were performed according to protocols approved by the Animal Ethics Committee of the Regierungspraesidium Giessen (permit numbers: G7/2017– No.844-GP and G11/2019–No. 931-GP).

## Consent for publication

All authors reviewed the results and contributed to the final manuscript. All authors approved this manuscript for publication.

## Availability of data and material

The scRNA-seq data are currently been deposited in GEO (accession number GSE pending). Genearrays data have been been deposited in GEO (accession number GSE162588).

## Competing interests

All the authors declare no competing interest

## Funding

S.B. was supported by grants from the Deutsche Forschungsgemeinschaft (DFG; BE4443/1-1, BE4443/4-1, BE4443/6-1, KFO309 P7, 284237345 and SFB1213-projects A02 and A04), UKGM, Universities of Giessen and Marburg Lung Center (UGMLC), DZL. J.S.Z was funded through a start-up package from Wenzhou Medical University and the National Natural Science Foundation of China (grant number 81472601). S.H. was supported by the UKGM (FOKOOPV), the DZL and University Hospital Giessen and grants from the DFG (KFO309 P2/8, 284237345; SFB1021 C05, SFB TR84 B9). DAA acknowledges support from NHLBI (R01HL141856). N.A. was funded through a start-up grant from the Cardio-Pulmonary Institute.

## Authors’contribution

N.A. designed the study, carried out the experiments, analyzed the data and wrote the manuscript. AL contributed to the experiments and quantification analysis. F.K. contributed to performing experiments, data analysis and writing of the manuscript. A.I.V.A. contributed to the experiments and quantification analysis. S.R contributed to the experiments and provided feedback in the writing of the manuscript. J.W. and J.K. contributed to the experiments and data analysis. S.H., G.B., J.Z, C.S. and D.A.A. provided feedback, helped shape the research, discussed the results, and contributed to the final manuscript. S.B. designed the project, regularly monitored the generated results, interpreted the results and wrote the manuscript in coordination with N.A. All authors reviewed the results and contributed to the final manuscript.

## Significance of the work

We demonstrate that Fgfr2b signaling is essential for alveolar epithelial lineage homeostasis in the adult mouse lung. Mature AT2s require Fgfr2b signaling for the maintenance of their proliferative capacity. The recently described injury activated alveolar progenitors (IAAPs) proliferate in the context of Fgfr2b deletion and functionally replace the loss of mature AT2s.

## Acknowledgements

We thank Stefan Guenther (Bioinformatics and deep sequencing platform at the Max Planck Institute for Heart and Lung) for help in ATAC-seq data analysis. We also thank Kerstin Goth for the animal husbandry and genotyping of the mice.

## Supplementary Figure captions

**Fig. S1 Fgfr2b is haplosufficient in AT2s.** FACS-based approach based on Pd-l1 expression to isolate AT2s and IAAPs in *Fgfr2b^+/+^* and *Fgfr2b^+/-^* lungs

**Fig. S2 Recombination efficiency in the IAAPs and AT2s in Exp. vs Ctrl lungs. a)** Ctrl and Exp. lungs were analyzed 36 hours after a single dose of Tam IP. FACS analysis was carried out to quantify the abundance of IAAPs and AT2s (out of Epcam) in Ctrl and Exp. lungs. **b)** Recombination efficiency in one-week and two-week tamoxifen treated animals. Quantification of the % of Tom^Pos^SftpcPos/SftpcPos by IF

**Fig. S3 Sequencing of the mutant transcript indicates complete deletion of *exon 8* in E-IAAPs isolated at day 7 during tamoxifen water treatment**

**Fig. S4 ATAC-seq analysis of E-IAAPs and C-IAAPs suggest that IAAPs get activated upon *Fgfr2b* deletion. a**) Coverage heat maps of C-IAAPs and E-IAAPs, displaying genome-wide regions of differential open chromatin peaks in E-IAAPs vs C-IAAPs. C-IAAP chromatin is less open and transcriptionally less active compared to E-IAAPs. ATAC-seq analysis of peaks based on the cutoffs shows 56 up-regulated in C-IAAPs (FDR < 0.05, l log_2_(FC) > 0.585, base Mean > 20), 455 up-regulated in E-IAAPs (FDR < 0.05, log_2_(FC) > 0.585, base Mean > 20) and 455 non-regulated (base Mean > 20, FDR > 0.5, log_2_(FC) between −0.15 and 0.15) which means 7.1% and 35.6% of the genome is differently accessible in C-IAAPs and E-IAAPs, respectively. **b**) Analysis of peaks obtained in the ATAC-seq experiment for E-IAAPs and C-IAAPs using Kobas for the Reactome database. Peaks overlapping gene body or near the transcription starting site of genes were annotated to the corresponding genes. All annotated peaks were split into lists of genes that display more open chromatin in E-IAAPs or C-IAAPs using DESeq2 on unified peak regions. Observed significance was adjusted by Benjamini-Hochberg correction for multiple tests (FDR). The resulting lists were used as input for Kobas to search for enriched terms in different databases. Top 7 terms were chosen by significance (FDR < 0.2). Results indicate that the term Metabolism is highly enriched in E-IAAPs, indicating that the chromatin of E-IAAPs is more accessible in loci of genes (gene body or promoter) associated with metabolism. Higher accessibility is associated with more transcriptional activity. Numbers in brackets display the number of identified genes / total number of genes for term in the database. DEG: Differentially expressed genes. Between brackets []: Genes found/total genes in term.

**Fig. S5 Viability of IAAPs and AT2s in Ctrl and *Fgfr2b-*cKO lungs following FACS.** Quantification of live and dead cells of Tom^Low^ and Tom^High^ by NucleoCounter, following FACS isolation from Exp. compared to Ctrl mice (n=4). Data are presented as mean values ± SEM. p < 0.05, p < 0.01, p < 0.001.

**Fig. S6 Analysis of the AT2s and IAAPs in Ctrl and Exp. lung in one-week tamoxifen followed by a two-week chase period a)** RT-PCR for detecting WT and *Fgfr2b* mutant transcripts in FACS-based sorted C-IAAPs, C-AT2s, E-IAAPs and E-AT2s. **b)** Flow cytometry analysis indicating expansion of the E-IAAPs as well as a global decrease in tdTom+ cells/Epcam+ in Exp. lungs. **c)** IF for tdTom+, Sftpc+ single positive cells as well as tdTom+ Sftpc+ double-positive cells.

**Fig. S7 Analysis of the AT2s and IAAPs in Ctrl and Exp. lungs in one-week tamoxifen followed by an eight-week chase period. a)** IF for Edu and TUNEL indicating a trend (non-significant) towards a residual increase in proliferation and apoptosis. **b)** Flow cytometry analysis indicating that the percentile of E-IAAPs, even though still higher than the one observed for C-IAAPs, is trending towards a normalization. Note that there is no change in the number of tdTom+/Epcam+ in Exp. and Ctrl lungs at this time point. **c)** IF for tdTom+, Sftpc+ single positive cells as well as tdTom+ Sftpc+ double-positive cells show no difference between Ctrl and Exp. lungs.

**Fig. S8 Enrichment of Pd-l1 expression in E-IAAPs vs E-AT2s. a)** Gene array for E-AT2s and E-IAAPs collected on day 4, 7 and 14 during the tamoxifen treatment. Note that *Cd33*, *Cd300lf* and *Cd274* (aka Pd-l1) are increased in E-IAAPs vs E-AT2s. **b)** Validation of these results by qPCR, cytospin and flow cytometry. **c)** qPCR indicates that *Cd33* and *Pd-l1* are enriched in E-IAAPs. **d)** Cystospin followed by IF for Pd-l1 indicates enrichment in Pd-l1 protein expression in E-IAAPs. **e)** Flow cytometry for E-IAAPs and E-AT2s followed by detection of Pd-l1 confirms that Pd-l1 is expressed chiefly in E-IAAPs.

**Supp figure for reviewers only.** Dynamic changes of IAAPs and AT2s during fibrosis formation and resolution. a) Two month-old SftpcCreERT2/^+^; TdTomato^flox/flox^ mice were treated with Tam IP (3 consecutive injections) and following a chase period of 7 days, were treated with saline or bleomycin. Mice were analyzed by flow cytometry to quantify the number of IAAPs and AT2s over total Tomato at day 10, 14, 16, 21, 28 and 60. b) Saline lung display the previously reported AT2 and IAAP ratio over tomato. c) quantification of the AT2s and IAAPs at the different time points. Representative flow cytometry FACS plot. d) Graph summarizing the dynamic changing in the ratio of IAAPs and AT2s during fibrosis formation and resolution, AT2s and IAAPs evolve in opposite direction.

## Notes

### Competing Interest Statement

The authors have declared no competing interest.

