## Supplementary figures and images for "Fgfr2b signaling is essential for the maintenance of the alveolar epithelial type 2 lineage during lung homeostasis in mice"

### supplementary data for Ahmadvand et al

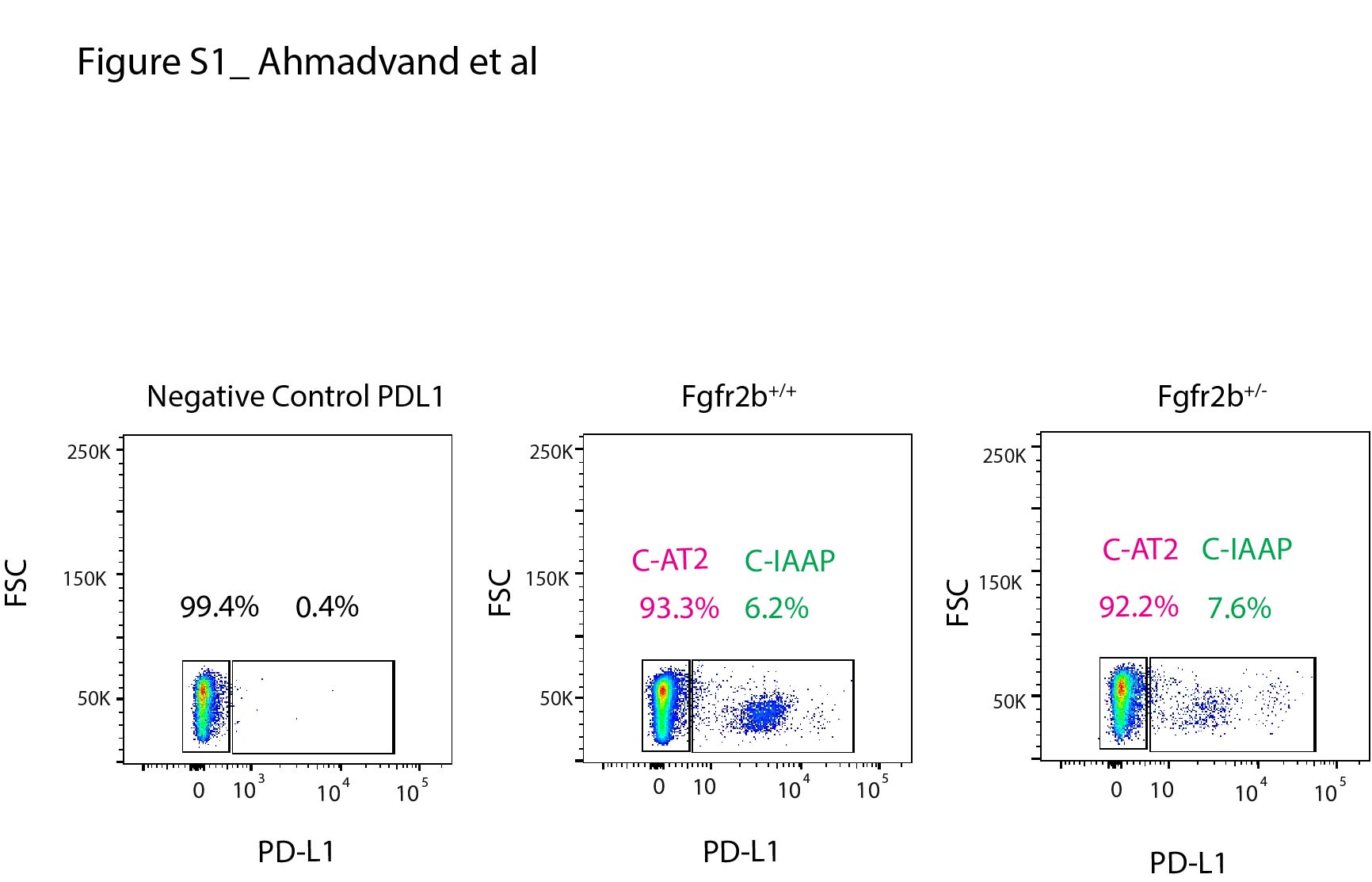

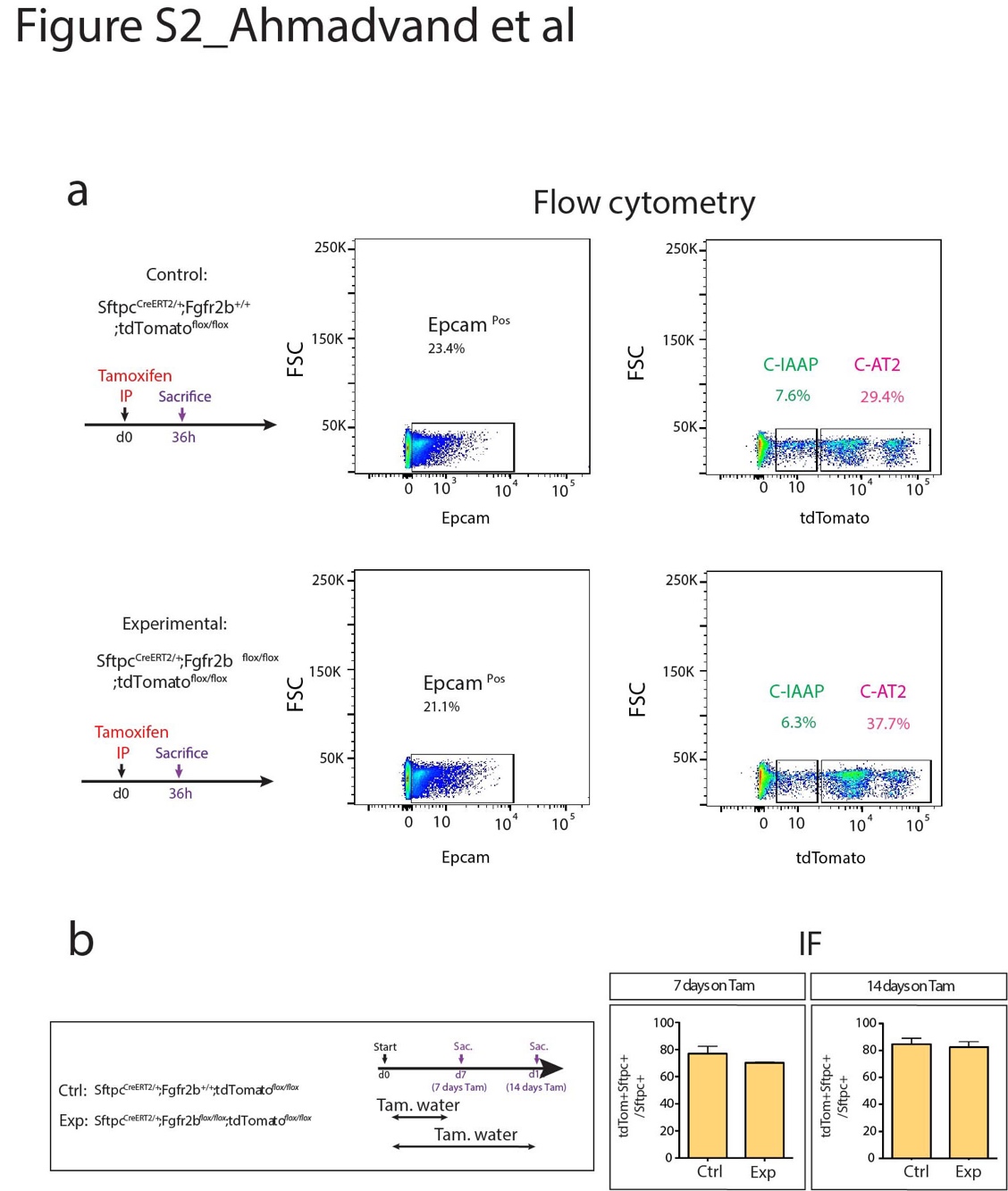

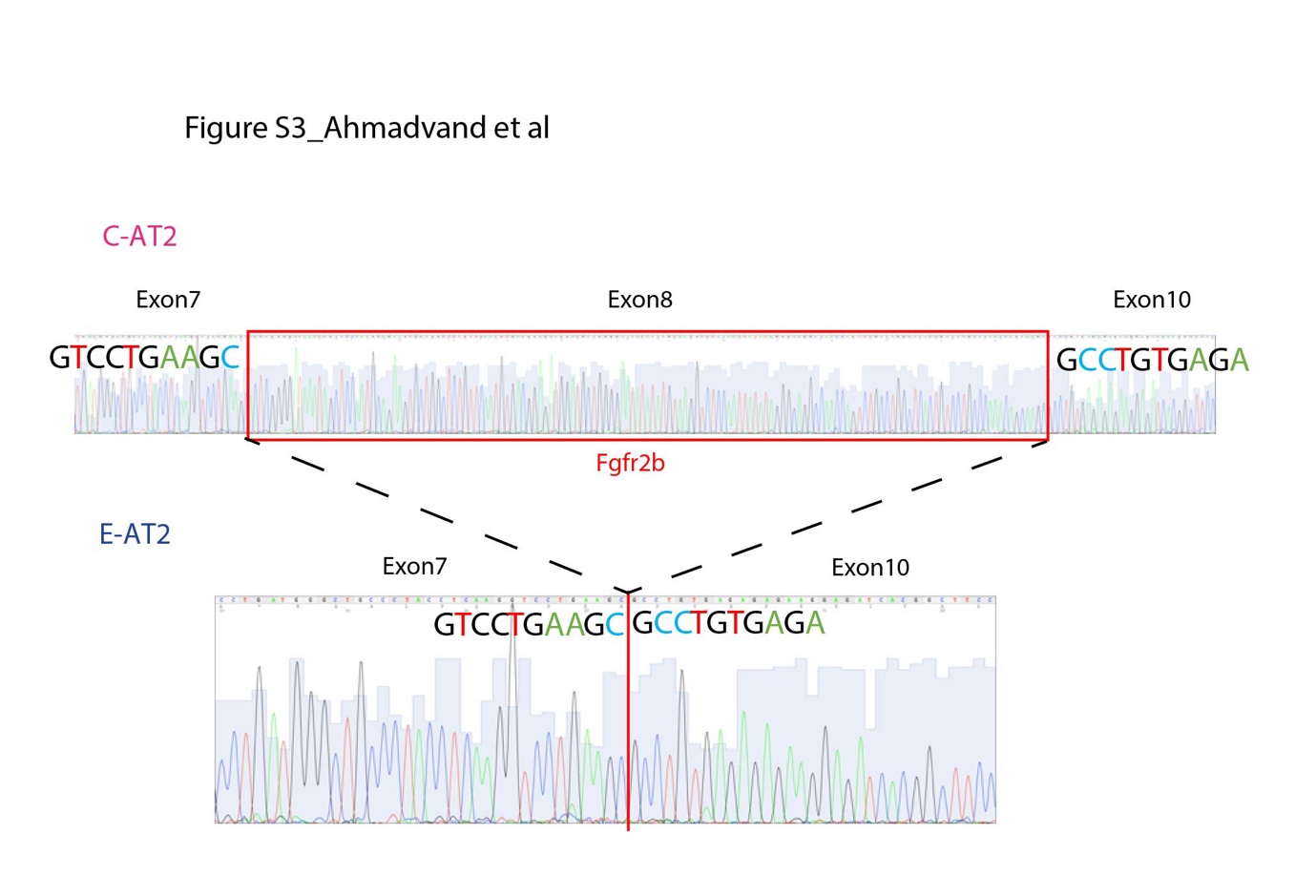

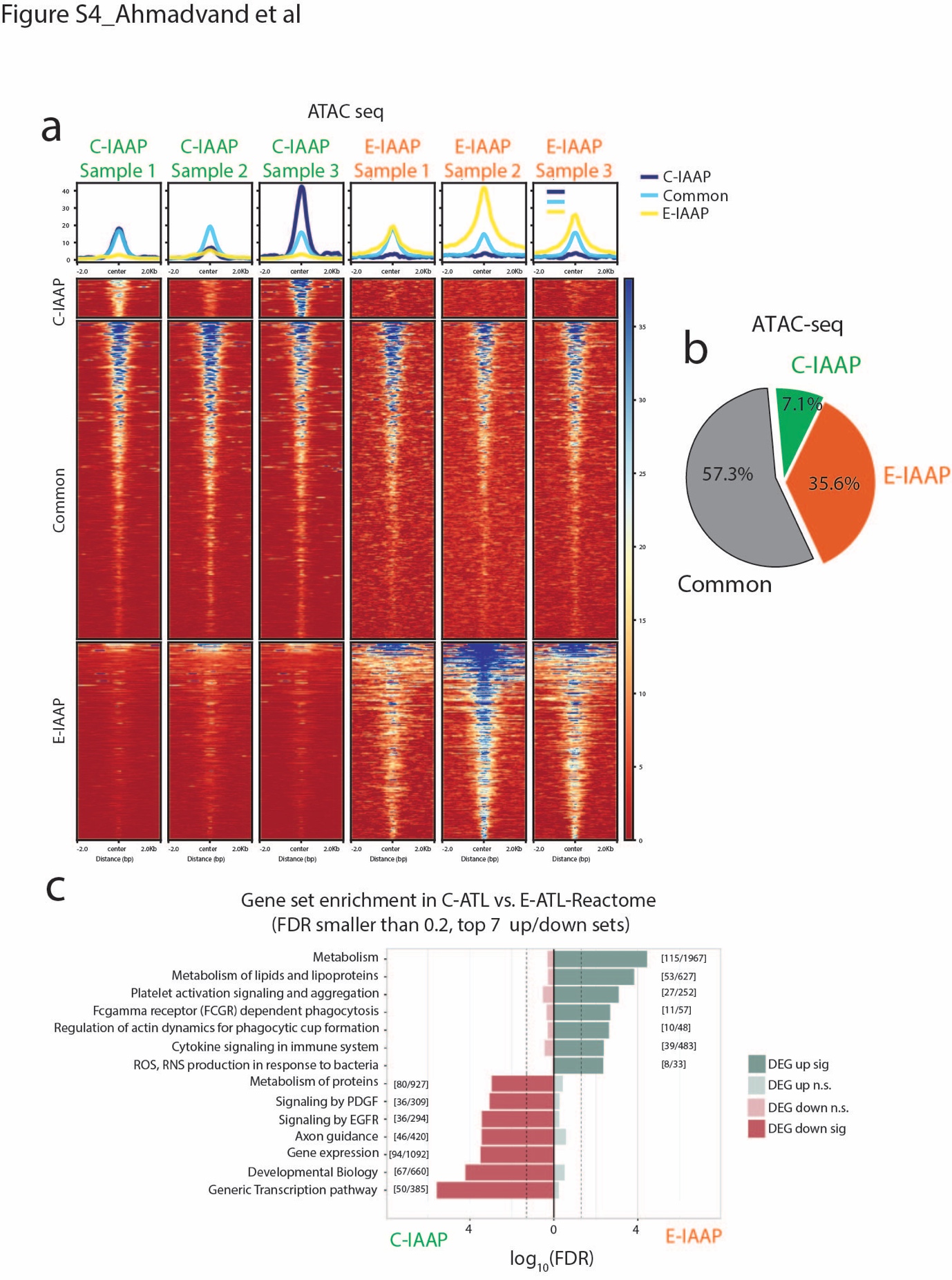

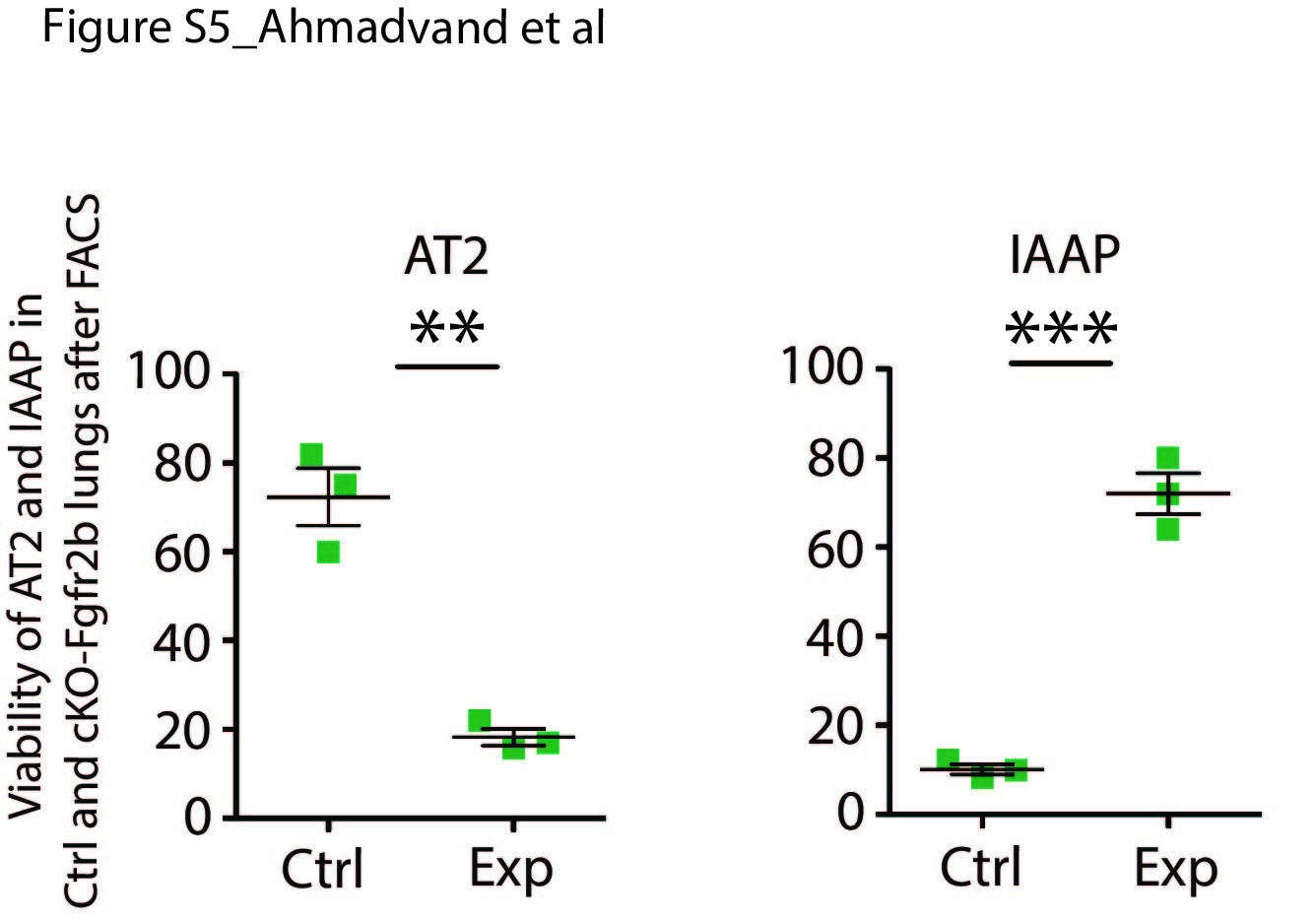

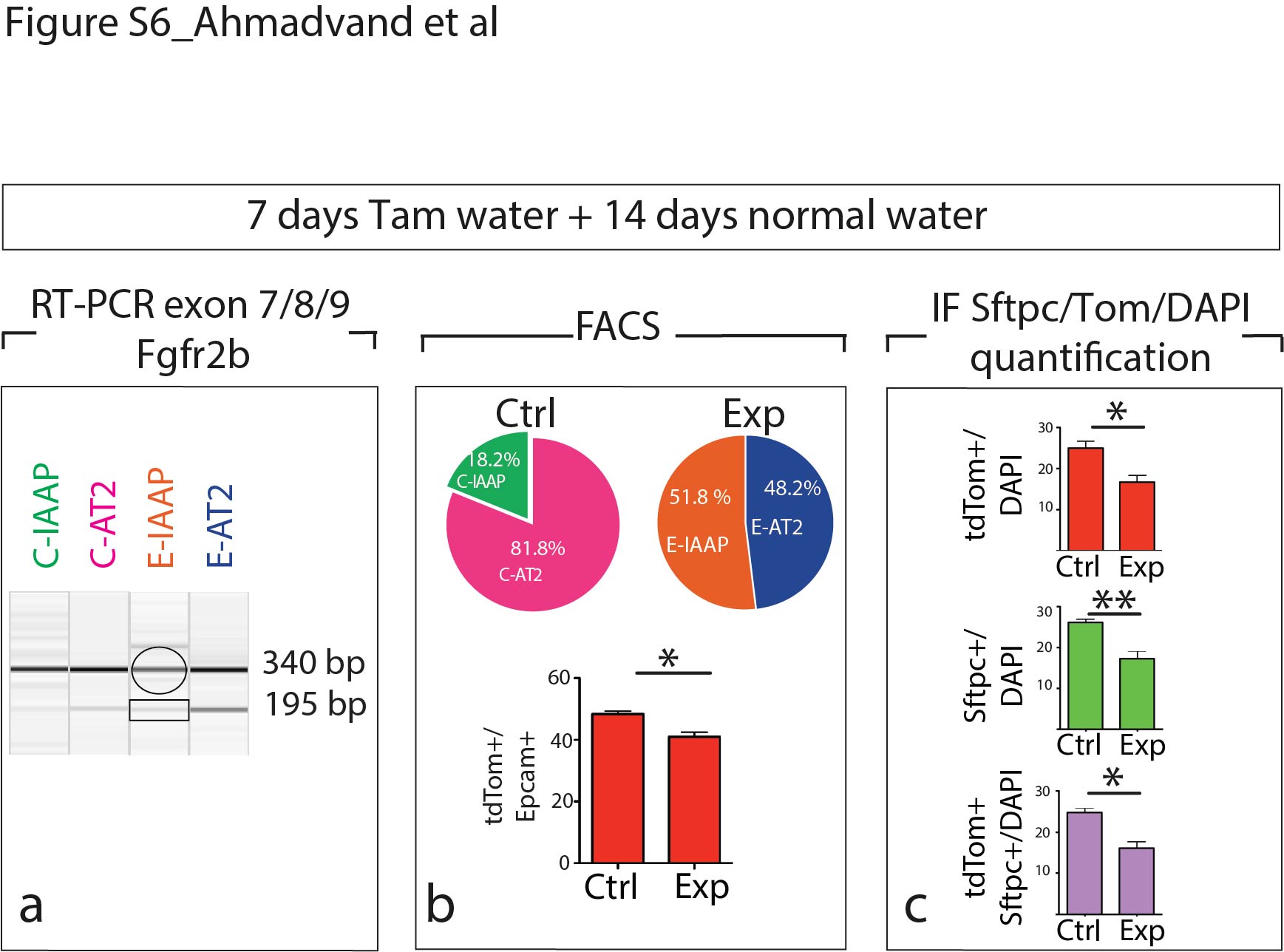

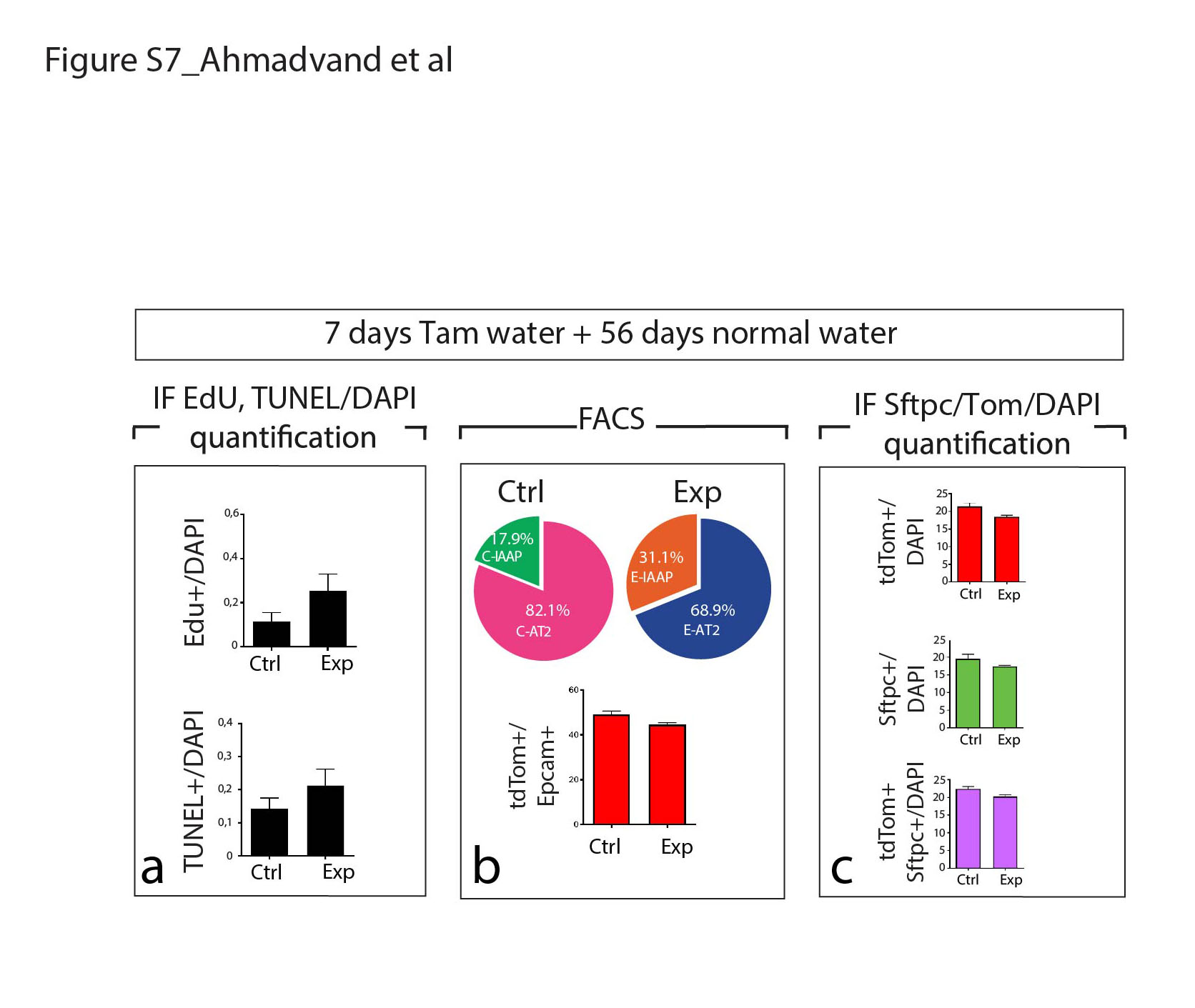

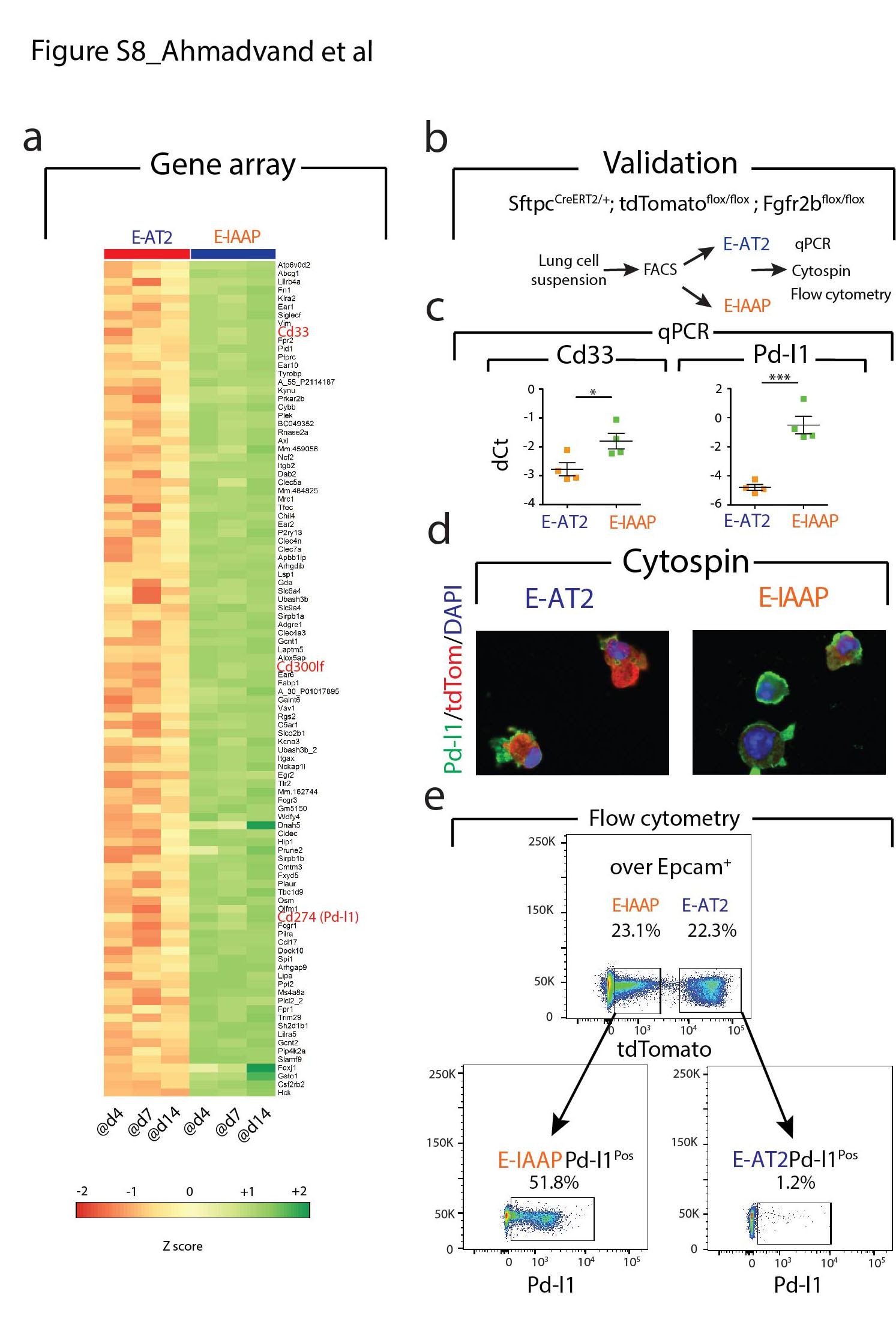
